# MODULARITY FACILITATES FLEXIBLE TUNING OF PLASTIC AND EVOLUTIONARY GENE EXPRESSION RESPONSES DURING EARLY DIVERGENCE

**DOI:** 10.1101/179424

**Authors:** Hannu Mäkinen, Tiina Sävilammi, Spiros Papakostas, Erica Leder, Leif Asbjørn Vøllestad, Craig Primmer

## Abstract

Gene expression changes have been recognized as important drivers of adaptation to changing environmental conditions. Little is known about the relative roles of plastic and evolutionary responses in complex gene expression networks during the early stages of divergence. Large gene expression data sets coupled with *in silico* methods for identifying co-expressed modules now enable systems genetics approaches also in non-model species for better understanding of gene expression responses during early divergence. Here, we combined gene co-expression analyses with population genetics to separate plastic and population (evolutionary) effects in expression networks using small salmonid populations as a model system. We show that plastic and population effects were highly variable among the six identified modules and that the plastic effects explained larger proportion of the total eigengene expression than population effects. A more detailed analysis of the population effects using a *Q*_*ST*_ - *F*_*ST*_ comparison across 16622 annotated transcripts revealed that gene expression followed neutral expectations within modules and at the global level. Furthermore, two modules showed enrichment for genes coding for early developmental traits that have been previously identified as important phenotypic traits in thermal responses in the same model system indicating that co-expression analysis can capture expression patterns underlying ecologically important traits. We suggest that module-specific responses may facilitate the flexible tuning of expression levels to local thermal conditions. Overall, our study indicates that plasticity and neutral evolution are the main drivers of gene expression variance in the early stages of thermal adaptation in this system.

## 1. Introduction

The relative roles of plasticity and evolutionary adaptation have gained considerable interest in recent evolutionary genetics research (Gienapp et al. 2008; Chevin et al. 2010; Merilä 2012; Crozier and Hutchings 2014; Merilä and Hendry 2014; Reusch 2014; DeBiasse and Kelly 2016). This is also tightly associated with a fundamental understanding of how populations adapt to rapid environmental changes (Franks and Hoffmann 2012). Rapid thermal adaptation may play a crucial role in future population persistence, particularly for ectotherms living in isolated habitats and thus unable to migrate to suitable thermal conditions (Franks and Hoffmann 2012; Narum et al. 2013). Rapid ecological responses to rising temperatures have been documented for several species, but the genetic mechanisms underlying these responses remain relatively poorly understood (Gienapp et al. 2008; Shaw and Etterson 2012; Merilä and Hendry 2014). In particular, the relative roles of plastic and evolutionary components underlying rapid ecological responses have remained challenging to demonstrate (Gienapp et al. 2008; Merilä 2012; Merilä and Hendry 2014). Plasticity, commonly understood as a capacity of the same genotype to express alternative phenotypes within the same generation, is widely acknowledged to produce rapid responses to new environmental conditions (Price et al. 2003; Crispo 2007; Fusco and Minelli 2010; Forsman 2015). Rapid genetic evolution in few generations has been demonstrated in a variety of model systems, challenging the traditional view of evolution as a slow process (Messer et al. 2016). Thus, there is potential for both processes to underlie rapid responses to abrupt environmental changes. According to current views, plasticity and evolution are not mutually exclusive, but they may interact during the adaptation to a new environment (Ghalambor *et al.* 2007; Ehrenreich & Pfennig 2015).

Several scenarios have been proposed to explain how plasticity and evolutionary responses might evolve during the course of adaptation (Pigliucci 2006; Crispo 2007; Schlichting and Wund 2014; Ehrenreich and Pfennig 2015; Hendry 2016). The initial response under novel environmental conditions might involve only a plastic response. Modelling, empirical and conceptual studies suggest that phenotypic plasticity enhances fitness, thus providing the capacity for survival (DeWitt et al. 1998; Price et al. 2003; Chevin et al. 2010; Fierst 2011; Draghi and Whitlock 2012; Lande 2015; Murren et al. 2015; Hendry 2016). Phenotypic plasticity may result in a nearly optimal phenotype, which is subsequently refined through natural selection when there is genetic variation in the same direction as the plastic response. As a result of this process, traits may become constitutively expressed, a phenomenon commonly known as genetic assimilation, or the environmental sensitivity may be maintained or increased (Baldwin effect) (Pigliucci 2006; Crispo 2007; Schlichting and Wund 2014; Ehrenreich and Pfennig 2015; Friedrich and Meyer 2016). Genetic assimilation may evolve when plasticity is, for example, costly and subsequent constitutive expression is favoured, whereas the Baldwin effect may be favoured under conditions in which maintaining plasticity is beneficial (Crispo 2007; Schlichting and Wund 2014). For example, environmental heterogeneity favours plasticity when the environmental cue is predictable (DeWitt et al. 1998; Crispo 2007; Hendry 2016). Furthermore, if plasticity drives the population to a new optimum in the new environment, then genotypes may also be shielded from natural selection, thereby constraining or slowing genetic evolution (Price et al. 2003; Ghalambor et al. 2007; Friedrich and Meyer 2016; Hendry 2016). Plasticity may also promote genetic evolution when the plastic response is maladaptive for favouring genetic compensation, known as counter-gradient variation (Conover and Schultz 1995; Morris and Rogers 2013; Hendry 2016).

Gene expression and its regulation is one of the key molecular mechanisms underlying plastic and evolutionary responses (Whitehead and Crawford 2006; López-Maury et al. 2008; Romero et al. 2012; Alvarez et al. 2014; DeBiasse and Kelly 2016). Epigenetic regulation via environmental stimuli may trigger plastic responses, whereas an evolutionary response may involve changes in regulatory elements (Hoekstra and Coyne 2007). Gene expression can show considerable flexibility when organisms are exposed to environmental gradients within the same generation, but it is also involved in long-term adaptation (López-Maury et al. 2008). Gene expression plasticity can be estimated using a genomic reaction norm approach by exposing populations to environmental variables in experimental settings. The slope of the genomic reaction norm can be informative about the magnitude of plasticity, and the genotype-environment interaction indicates genetic variation in plasticity (Aubin-Horth and Renn 2009). Estimating evolutionary responses in gene expression remains a challenge, reflecting the lack of an appropriate null model for separating variance as a result of neutral divergence from natural selection (Fraser 2011; Harrison et al. 2012; DeBiasse and Kelly 2016). Furthermore, methods using phylogenetic relationships to infer selection in expression data might not be applicable to closely related populations (Rohlfs et al. 2014). Q_ST_-F_ST_ (or P_ST_ for phenotypic data) comparisons are widely used to infer local adaptation in phenotypic traits but have also been applied relatively rarely to ‘omics’ data (Leinonen et al. 2013). In this approach, the inference of adaptive evolution is based on the presumably neutral distribution of the F_ST_ estimated from genetic markers to which the distribution of Q_ST_ is contrasted. The Q_ST_ estimates outside the F_ST_ distribution are putative candidates for natural selection (Leinonen et al. 2013). Modern sequencing technologies enable the simultaneous collection of gene expression and genetic variation data for a large number of molecular phenotypes, providing a meaningful starting point for estimating the evolutionary forces affecting expression divergence (De Wit et al. 2015).

Similarly, large gene expression data sets coupled with *in silico* methods for identifying co-expressed gene networks or modules enable a systems genetics approach, even in non model species (Soyer and O’Malley 2013; Feltus 2014). Analyses of co-expressed gene networks have been widely used in medical genetics but are also gaining popularity in evolutionary genetics (Langfelder and Horvath 2008; Feltus 2014; Ruprecht et al. 2017). The rationale behind *in silico* co-expression gene network analysis is that gene expression correlation may reveal functionally related genes belonging to the same biological pathway (Langfelder and Horvath 2007; Langfelder and Horvath 2008). Furthermore, the expression variance of genes belonging to a module can be summarized to eigengenes, and their expression can be further analysed in relation to external information. Thus, multiple testing problems can be reduced compared to testing each gene separately to detect differential expression (Langfelder and Horvath 2008). Genes and gene products interact in complex networks. The position of a gene in a network or the number of interactions to other genes can affect the evolutionary dynamics of gene expression (Levy and Siegal 2008; Feltus 2014; Fischer et al. 2016; Laarits et al. 2016). For example, the number of protein-protein interactions and the location of the gene in a network may constrain or buffer against changes in gene expression (Han et al. 2004; Levy and Siegal 2008; Papakostas et al. 2014). Thus, analysing gene expression changes within and among networks can provide further insights into how populations have adapted to local conditions (Ruprecht et al. 2017).

Understanding the roles of plastic and evolutionary responses in gene expression may benefit from integrating methods commonly used for evolutionary and systems biology (Soyer and O’Malley 2013; Feltus 2014). Co-expression analysis might reveal complex interaction networks but are not informative of the evolutionary forces shaping the network evolution (Soyer and O’Malley 2013). Traditional Q_ST_-F_ST_ comparisons might help separate neutral and adaptive processes in network evolution and provide a global view of transcriptome divergence. Comparisons of recently diverged populations may provide further insights into the molecular mechanisms underlying rapid thermal adaptation in a time scale comparable to anthropogenic environmental change. Furthermore, direct comparisons of ancestral-derived populations are informative of the evolution of plasticity (Schlichting and Wund 2014). Here, we used European grayling (*Thymallus thymallus*) populations inhabiting small mountain lakes in Norway to investigate early-stage divergence in gene expression. This model system is suitable for investigating the early stages of divergence because colonization dates back 25-30 generations (Haugen and Vøllestad 2000; Haugen and Vøllestad 2001). In addition, knowledge concerning the ancestral population facilitates comparisons to the derived populations, enabling the tracking of evolutionary sequences and plastic events (Schlichting and Wund 2014).

Here, we investigate the relative roles and the interactions of plastic and evolutionary responses in gene co-expression networks during rapid thermal adaptation. We focus on two working hypothesis. First, under the genetic assimilation scenario, the plastic response to thermal treatment is lost during the course of divergence, and the populations show divergent expression profiles, reflecting adaptive evolution. Second, under the Baldwin effect scenario, plasticity is maintained or even elevated relative to the ancestral level, but populations may also show divergence in gene expression. To address the abovementioned questions, we raised developing embryos from four grayling populations originating from varying thermal environments and exposed them to two thermal treatments in a common garden environment. To evaluate the above scenarios, we first used a gene co-expression analysis to identify expression modules of potentially functionally similar transcripts. We then analysed module eigengene expression variation in an ANOVA framework to partition variance to treatment, population and their interaction effects. ANOVA analysis should reveal the relative contributions of plastic (treatment) and population (evolutionary) effects on module eigengene expression. Second, we further analysed the population effect in gene expression using a broad sense Q_ST_-F_ST_ comparison. This approach is used to estimate gene expression variation resulting from neutral and potentially adaptive processes within modules and at the global level.

## Results

On average, 78.7 million paired-end reads were obtained per sequencing library. After quality filtering and removing PCR duplicates, 68.1 million reads remained (86.5%) (supplementary table 1). The average GC content of the quality-filtered libraries was 46%. *De novo* assembly with Trinity identified 142653 transcripts (including isoforms) and 109102 trinity ‘genes’. The total length of the de novo assembly was 143.923 Mb, and the mean and average contig lengths were 583 bp and 1009 bp, respectively. *In silico* prediction of the putative coding sequence with TransDecoder identified 136291 transcripts with open reading frames. Clustering of highly similar sequences with CD-hit identified 61190 unique proteins. The annotation of the unique proteins with reciprocal blast search identified 19461 putative homologies to at least one of the species (zebrafish, stickleback, cod or Atlantic salmon). 131 putative paralogous genes i.e. the same transcript showed reciprocal blast hits to different gene models were removed from the down stream analyses. Specifically, 6069 (31.2%), 3830 (19.7%), 4302 (22.1%) and 5260 (27.0%) transcripts had one, two, three or four BLASTP hits, respectively. Therefore, the majority of the transcripts (13392, 68.8%) had more than one reciprocal blast hit in reference species. Approximately 27.3 (40.1%) million reads were mapped back concordantly to the de novo assembly with Bowtie2 (supplementary table 1). Altogether count data were obtained from 19330 annotated transcripts. However, the final data set comprised 16622 annotated transcripts after the removal of transcripts containing at least one individual with zero counts. This filtering step was applied to remove transcripts showing uninformative signals and to avoid frequent crashes in the WGCNA resampling analysis.

**Table 1.** ANOVA results on the module eigengene expression variation as a result of population, treatment and their interaction effects.

| Module | Population (13, 27<sup>1</sup>) | Treatment (1, 27) | Interaction (3, 27) |
| --- | --- | --- | --- |
| black | 13.2<sup>2</sup> (2.076<sup>3</sup>) | 27.7 (13.036)*** | 1.6 (0.249) |
| blue | 21.0 (25.932)*** | 57.4 (212.624)*** | 14.4 (17.763)*** |
| brown | 1.8 (0.19) | 0.3 (0.108) | 14.5 (1.569) |
| green | 9.6 (1.615) | 35.7 (18.097)*** | 1.4 (0.228) |
| red | 0.1 (0.015) | 1.2 (0.358) | 11.6 (1.202) |
| turquoise | 23.6 (55.219)*** | 54.6 (383.606)*** | 18.0 (42.22)*** |

\*\*\* P < 0.001
<sup>1</sup>Degrees of freedom
<sup>2</sup>Percentage of variance explained
<sup>3</sup>F ratio

Principal component analysis showed differentiation both between treatments and populations (figure 3). PC1 explained 25.9% and PC2 explained 17.3% of the total variation in gene expression. There was clear differentiation between populations in cold and warm treatments for R. Gudbrandsdalslågen, L. Lesjaskogsvatnet and L. Aursjøen. However, L. Hårrtjønn in the cold treatment overlapped other populations in the warm treatment (figure 1). Analysis of variance using PC1 as a dependent variable revealed significant population [F(3, 27) = 26.82, P<0.001], treatment [F(1, 27)=208.12, P<0.001] and their interaction effects [F(3, 27) = 11.25, P<0.001]. Post hoc (Tukey HSD) tests indicated that three out of six pairwise population comparisons in the cold treatment were significant (adjusted p-value < 0.05), whereas in the warm treatment, two comparisons were significant (supplementary figure 3). When different populations were compared between the treatments only two out of sixteen comparisons were non-significant (supplementary figure 3).

**Figure 1.**
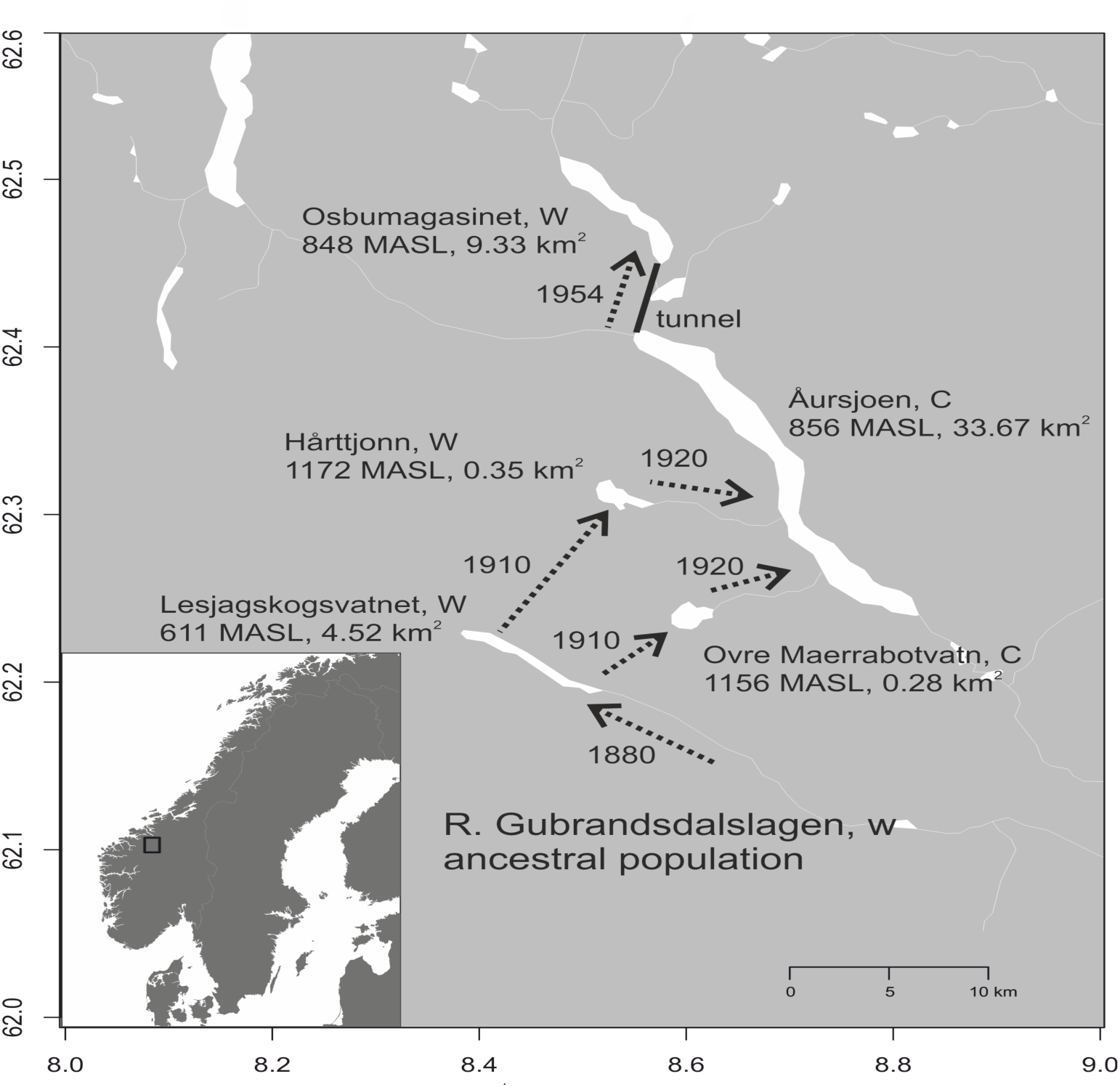
A map of the Grayling study system showing the colonization routes (arrows) and timing (numbers along the arrows) as inferred from the historical records, lake size (km^2^), and elevation (MASL=meters above sea level). These lakes differ in their thermal profiles during the grayling development period and can be roughly classified as cold (C) and warm (W) populations.

**Figure 2.**
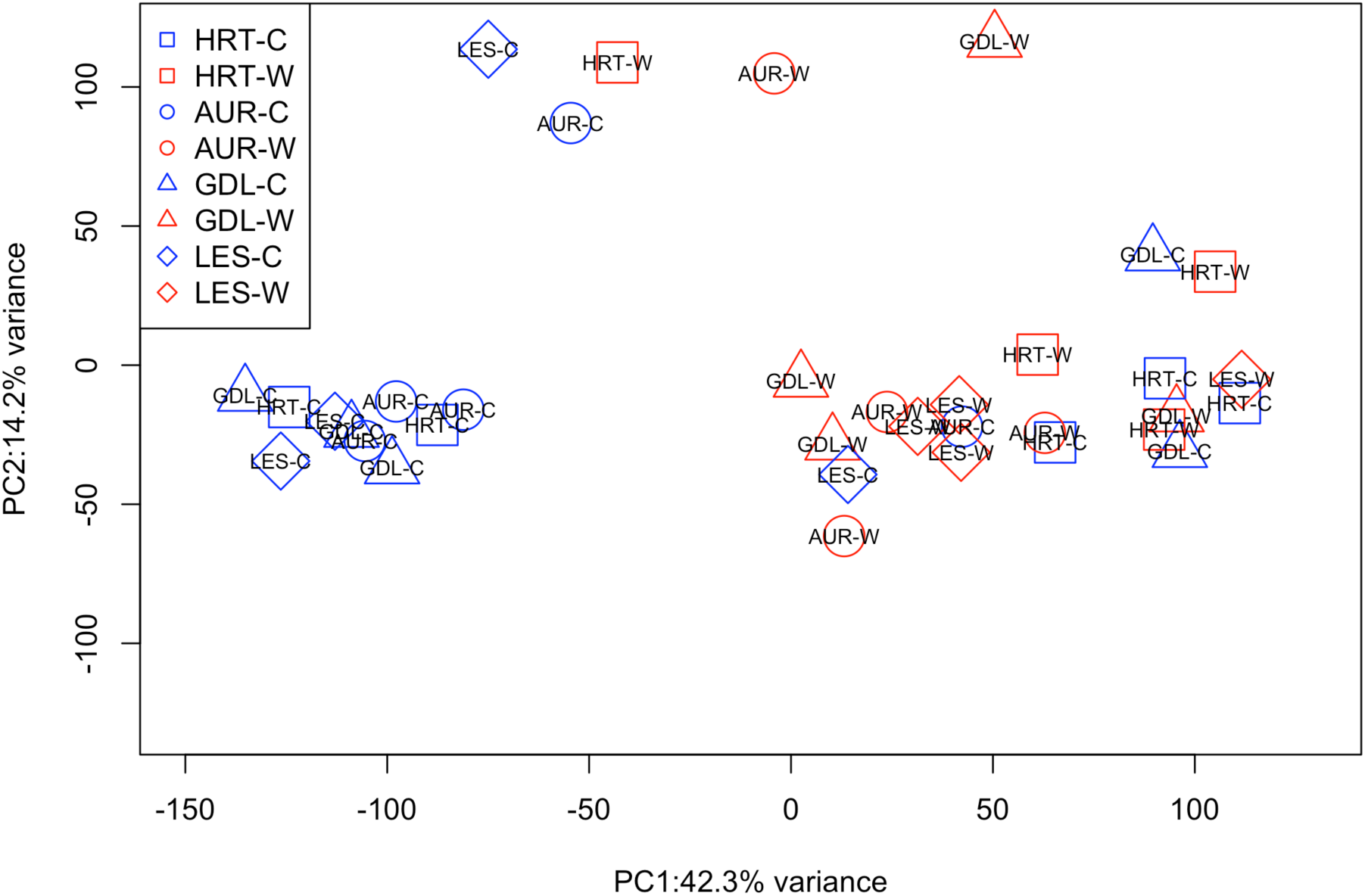
Principal component analysis (PCA) of the un-normalized gene expression data. GDL=Gudbrandsdalslågen, HRT=L. Hårrtjønn, AUR=L. Aursjøen and LES=L. Lesjagskogsvatnet. C indicates cold treatment and W indicates warm treatment.

**Figure 3.**
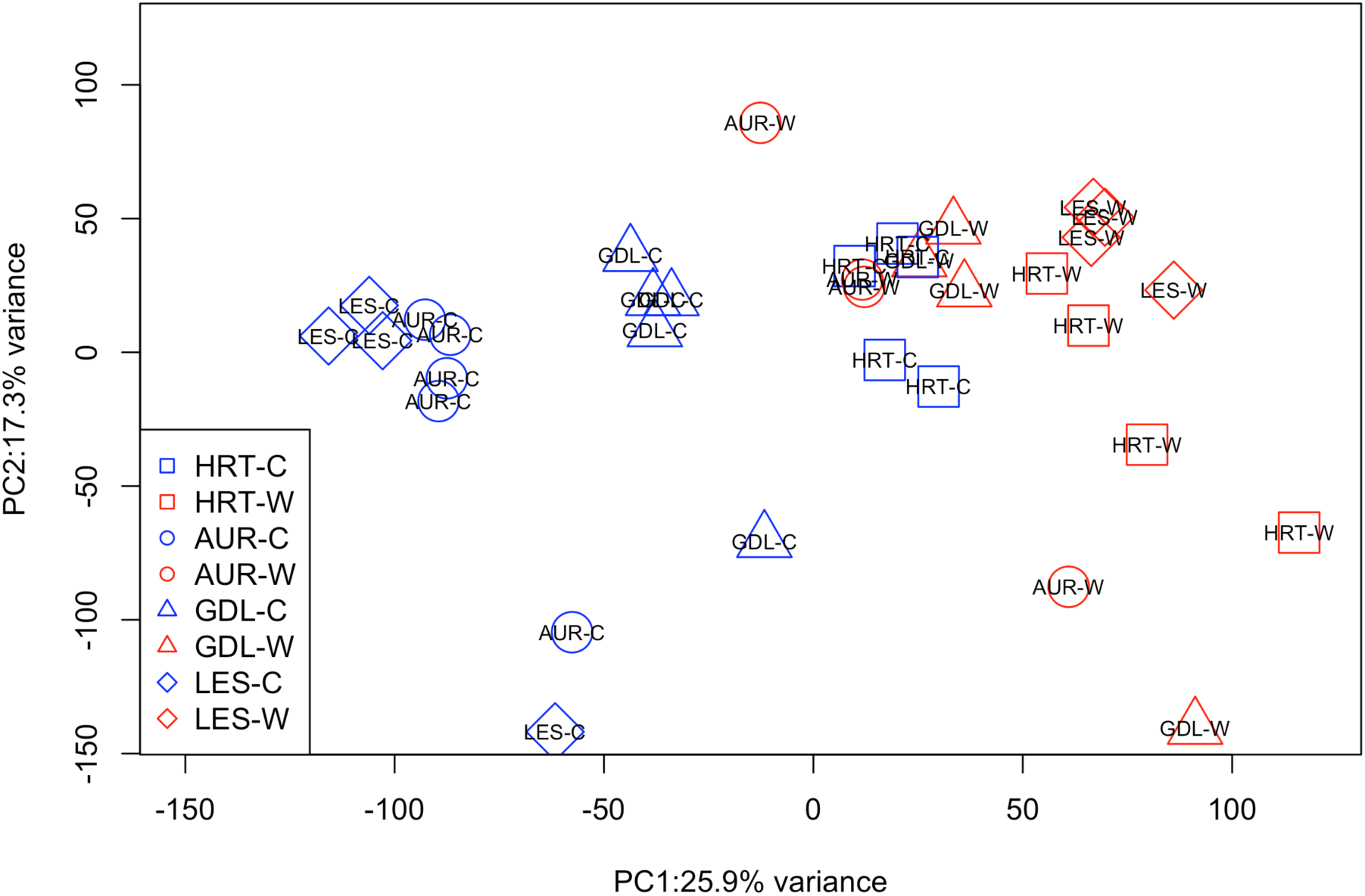
Principal component analysis (PCA) of the residual normalized gene expression data, showing the relationships among treatments and populations. GDL=Gudbrandsdalslågen, HRT=L. Hårrtjønn, AUR=L. Aursjøen and LES=L. Lesjagskogsvatnet. C indicates cold treatment and W indicates warm treatment.

The weighted gene co-expression analysis identified ten modules, and six modules (assigned to black, blue, brown, green, red and turquoise colours by the WGCNA analysis) were robust in the resampling analysis, i.e., showing significant overlap in 70% of the resampled data sets with the original data set (figure 4). The six statistically robust modules contained a total of 5999 (36.1%) transcripts. Black, blue, brown, green, red and turquoise modules contained 223, 1499, 1133, 740, 302 and 2102 transcripts, respectively. The majority of the transcripts (9496, 57.1%) were not assigned to any particular module (grey module) and the remaining modules (magenta, pink and yellow) showed instability in the re-sampling analysis (figure 4). PC1 on the transcripts belonging to the statistically robust modules explained 96.5% of the total variation, whereas the PC2 explained 1.4% of the total variation (figure 5). Module eigengenes (PC1) showed variable responses to population, treatment and their interaction effects (table 1, figure 6). Blue and turquoise modules had significant (P < 0.001) population, treatment and their interaction effects, whereas black and green modules showed only significant treatment effects (table 1, figure 6). The other modules (red and brown) showed non-significant population, treatment or their interaction effects (table 1, figure 6).

**Figure 4.**
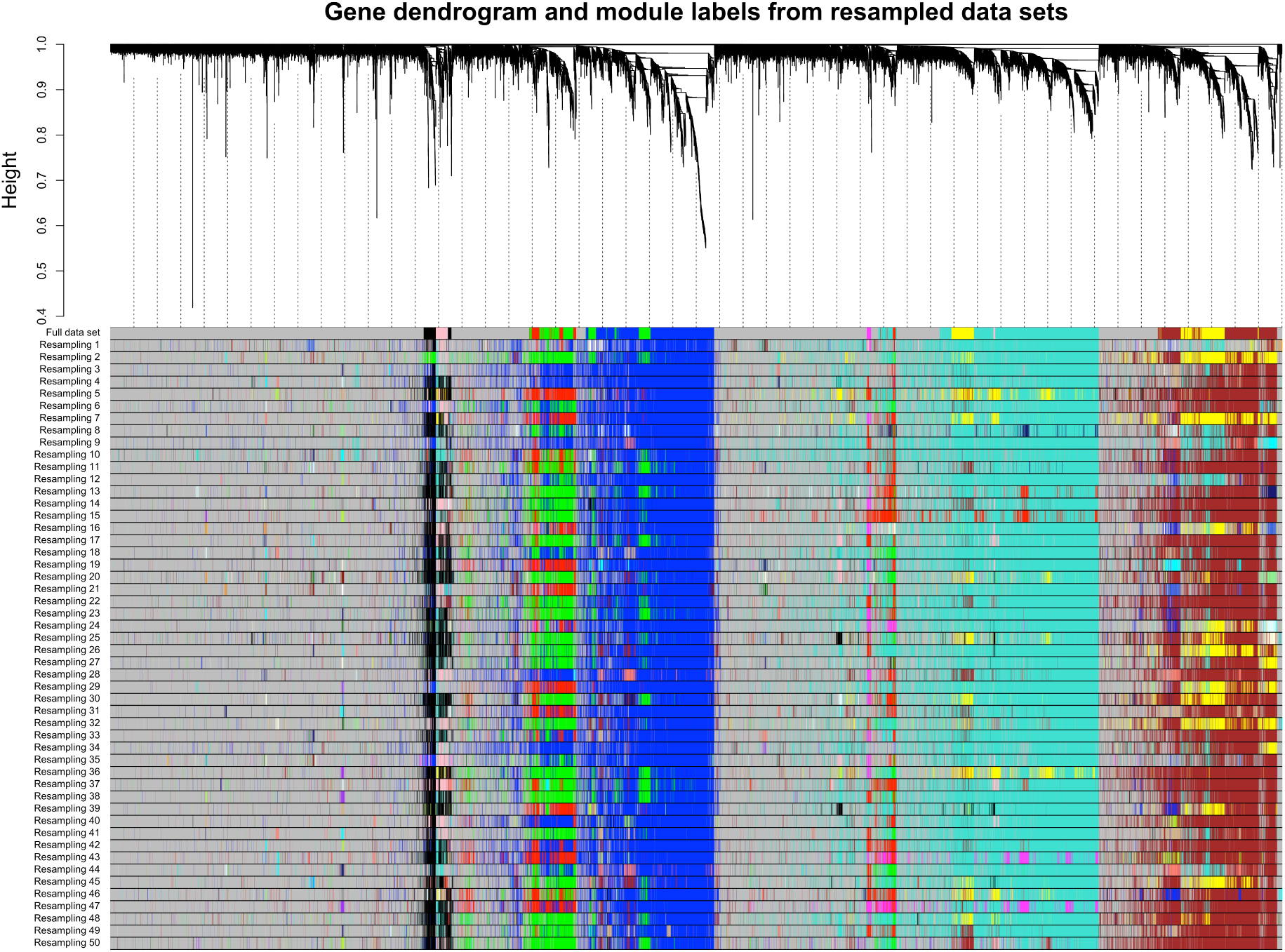
Results of the weighted gene co-expression analysis showing the dendrogram of transcripts based on the co-expression similarity. Each transcript is assigned to a module described with different colours. The stability of the modules was examined using one hundred re-sampling replicates, but the first fifty re-sampled data sets are shown for clarity.

**Figure 5.**
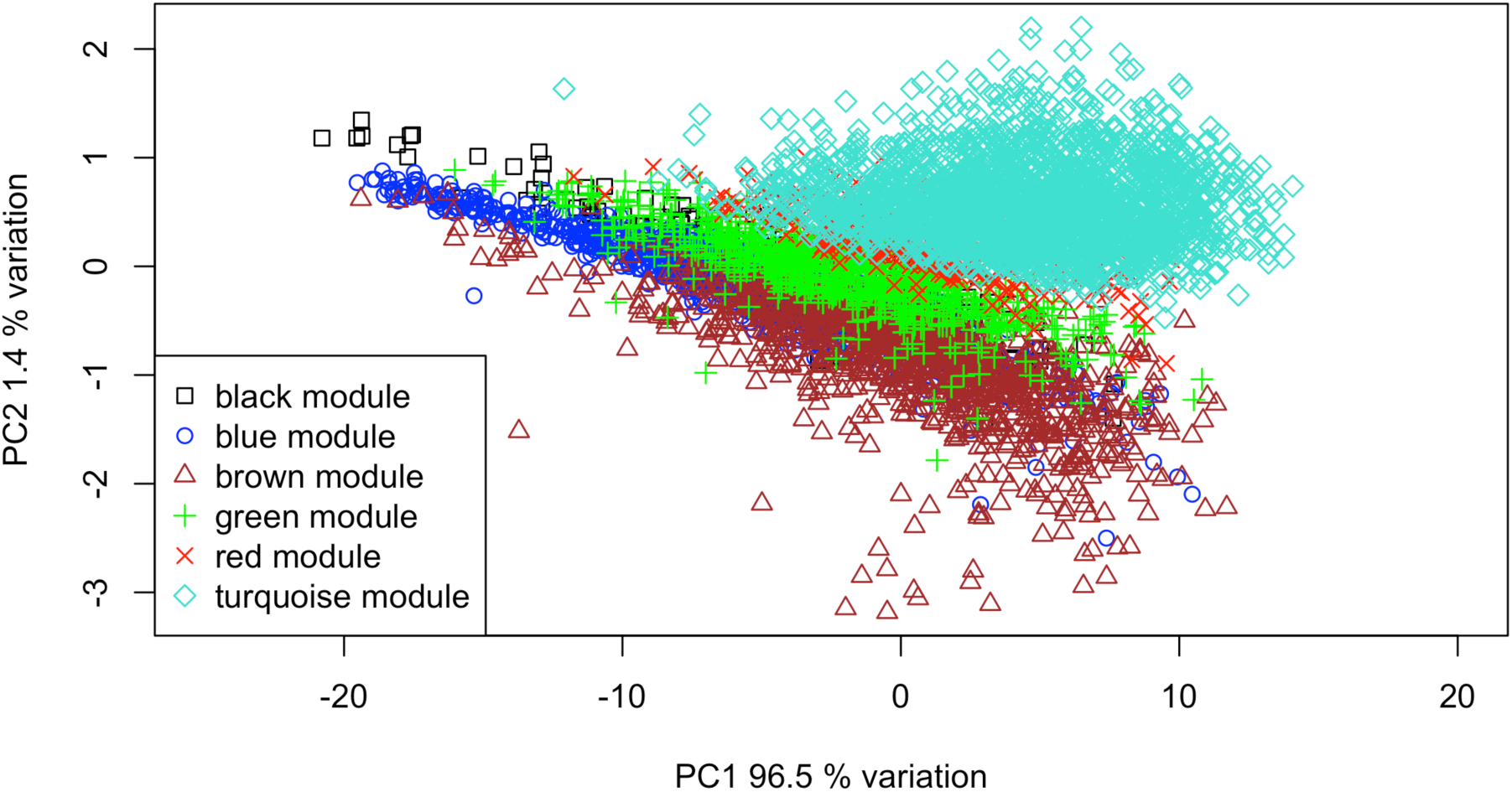
PCA on transcripts across all treatments and populations belonging to six statistically robust modules showing divergence in gene expression.

**Figure 6.**
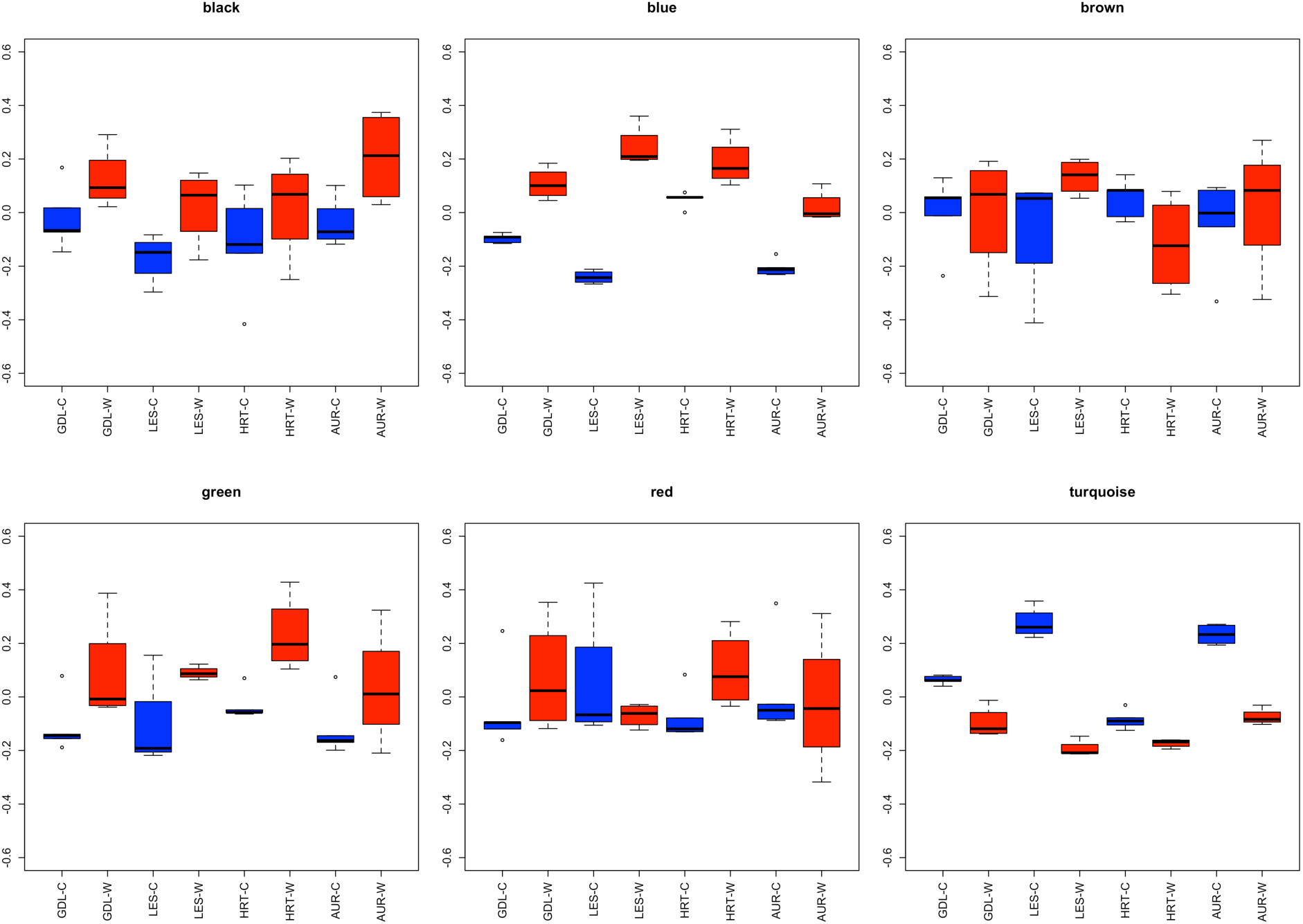
Module eigengene variation among the populations and treatments.

The mean and range of Q_ST_ across all transcripts were 0.024 (0-0.570), 0.044 (0-0.726) and 0.062 (0-0.799) assuming 0.5, 1 and 1.5 for the c/h^2^ scalar variable, respectively (supplementary figure 4). A total of 13121 transcripts were significant, i.e., the confidence intervals excluded zero. The mean F_ST_ across 2458 SNP loci was 0.128 and the range of locus specific F_ST_ values was 0-0.531 (supplementary figure 4). There were no differences in the mean F_ST_ in SNPs located in the flanking regions (F_ST_=0.129, n=1955), and non-synonymous positions (F_ST_=0.129, n=256) or synonymous positions (F_ST_=0.116, n=247). SNP data separated populations in PC1 (16.4% variation) and PC2 (11.8% variation), whereby the ancestral population was the most distant from the other populations (supplementary figure 5). The Q_ST_ estimates of two transcripts [TR19626: 0.799 (95% C.I. 0712-0.871) and TR47182: 0.709 (95% C.I. 0.576-0.813)] fell outside the upper range of the F_ST_ distribution when c/h^2^=1.5, and one transcript [TR19626: 0.726 (95% C.I. 0.622-0.819)] fell outside the upper range of the F_ST_ distribution when c/h^2^=1.0 (supplementary figure 4). No Q_ST_ estimates were detected outside the lower range of the F_ST_ distribution (supplementary figure 4). Thus, almost the entire range of Q_ST_ estimates fell within the F_ST_ distribution, indicating that the transcriptome divergence can be explained by patterns consistent with neutral evolution. However, the mean Q_ST_ differed between modules. Black (0.037), brown (0.020), green (0.041) and red (0.016) modules showed relatively small differentiation, whereas blue (0.123) and turquoise (0.137) modules showed a higher degree of differentiation on average assuming c/h^2^=1.5 (figure 7).

**Figure 7.**
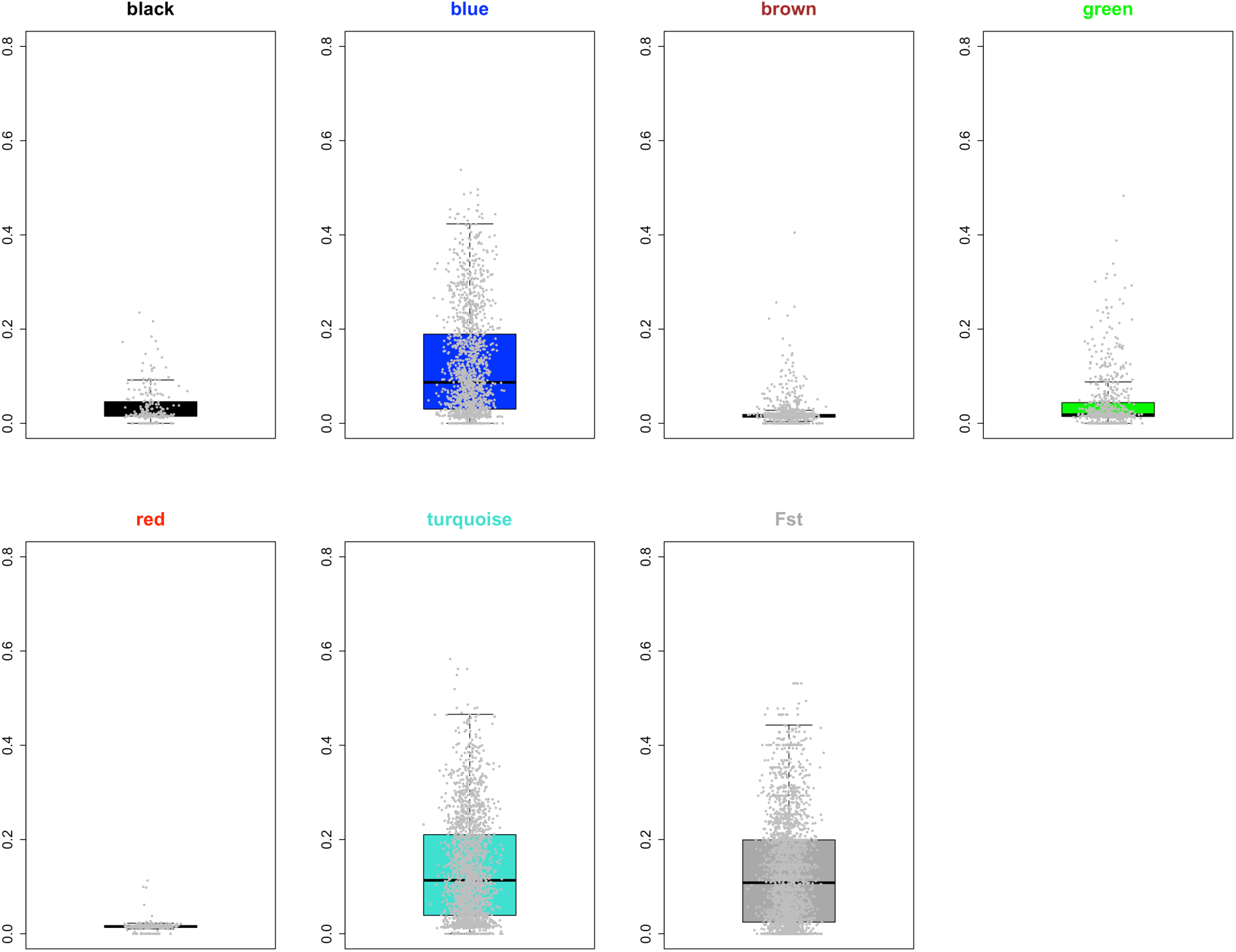
Boxplots depicting the variance of QST for each of the transcripts in each module, assuming scaling factor 1.5 (i.e., c/h^2^). The distribution of F_ST_ for 2458 SNP loci is shown for comparison.

Altogether 126 (56.5%), 1060 (70.7%), 665 (58.7%), 440 (59.5%), 165 (54.6%) and 1275 (60.7%) zebrafish annotations were recovered for the grayling transcripts belonging to the black, blue, brown, green, red and turquoise modules, respectively, which were used for enrichment analysis using the STRING database. Gene enrichment analysis revealed significant (FDR < 0.05) 33 (black module), 134 (blue module), 112 (green module), and 18 (turquoise module) GO terms associated with biological processes. After merging semantically similar categories, 15, 55, 33, and 7 categories remained for black (supplementary figure 6), blue (supplementary figure 7), green (supplementary figure 8) and turquoise (supplementary figure 9) modules, respectively. No significant enrichments for the biological processes were observed for the red and brown modules. Most of the categories belonged to general terms, such as “biological regulation” and “cellular process”, both of which were identified in all modules, except the turquoise module. Other categories were involved in biological functions “response to stress”, “response to stimulus” (green module) and “methylation” (blue module). There were also several more specific terms associated with developmental processes. The black module contained “muscle fibre development”, the green module “nervous system development” and the turquoise module “embryonic organ development” terms. Altogether six (blue module), two (brown module), two (green module), one (red module) and five (turquoise module) significant (FDR <0.05) Pfam protein domain enrichments were identified (supplementary table 2).

## Discussion

### The evolution plastic response during early divergence

One of the major findings of the present study is that populations respond differently to the thermal treatment at both the whole transcriptome level and within the transcriptional modules. This finding opens up possibilities for disentangling the causes of variable responses in the context of how plasticity itself evolves and how it interacts with evolutionary responses. Studies have suggested that the initial response to new environmental conditions is produced through plasticity, but if there is genetic variation in the same direction, then the response can become genetically determined. During this process, the environmental sensitivity or plasticity can be lost or reduced, a phenomenon known as genetic assimilation or accommodation (Pigliucci 2006; Crispo 2007; Schlichting and Wund 2014; Ehrenreich and Pfennig 2015; Friedrich and Meyer 2016). However, plasticity can be maintained or increased relative to ancestral plasticity levels, a phenomenon known as the Baldwin effect (Crispo 2007). It is expected that the Baldwin effect is favoured when plasticity is not costly and is beneficial. Genetic assimilation, however, is expected to evolve under conditions in which constitutive expression is favoured. The cost of plasticity and maladaptive response are expected to favour genetic assimilation (Crispo 2007). If the genetic assimilation scenario would hold in this study system, then we would expect a loss of plasticity and increased population effects during the colonization of different habitats. By contrast, we found that the levels of plasticity were not reduced but rather varied among populations and were not related to the level of ancestral plasticity. One point of speculation might be that genetic assimilation potentially takes longer than the divergence between the study populations (25-30 generations). Experimental studies have shown that genetic assimilation in thermal stress can rapidly evolve (ten generations) in laboratory populations of nematodes (Sikkink et al. 2014). In bacterial populations, genetic assimilation to high CO_2_ levels evolved after 4.5 years (570-850 generations) of experimental evolution (Walworth et al. 2016). Although the results from experimental evolution studies in other species cannot be directly translated to natural populations, there are indications that genetic assimilation can rapidly evolve. Although there is potential for rapid evolution of genetic assimilation, the response at the gene expression level can be complicated. Sikkink *et al.* (2014) showed that there were no correlated responses in gene expression to the evolved changes in thermal stress at the phenotypic level. This finding may further complicate detecting genetic assimilation at the molecular level.

Heterogeneous environments can favour plasticity because multiple optima are needed during the life-time of an organism (Crispo 2007; Murren et al. 2015; Hendry 2016), potentially explaining the variable plastic response observed in the grayling populations. We observed the lowest plastic response in the smallest lake (L. Hårrtjønn) with few river outlets. By contrast, we observed the large plastic responses in larger lakes with many small tributaries. For example, in L. Lesjaskogsvatnet, the grayling spawns in numerous ‘large cold’ and ‘small warm’ rivers, which differ in their thermal profiles (Haugen 2000; Haugen and Vøllestad 2001; Gregersen et al. 2008; Kavanagh et al. 2010). The embryos hatch, and the larvae drift or migrate from the spawning tributary into the lake during summer and early autumn. Grayling in L. Lesjaskogsvatnet may thus experience a wide thermal range during its lifetime, favouring larger plasticity. Similar patterns may arise if plasticity is costly in one environment but not in the other. Costs may arise as a result of energetic costs of maintaining genetic machinery for producing a plastic response, developmental instability and genetic costs if the plasticity is associated with a disadvantageous gene (DeWitt et al. 1998; Crispo 2007). Studies investigating plasticity costs have reported variable outcomes, but overall the results suggest that the costs are small or absent (Snell-Rood et al. 2010; Murren et al. 2015). Evaluating the costs of plasticity in grayling populations in the present study would be difficult, but the relation of enrichments to growth and developmental traits might indicate that plasticity costs could indeed exist. Kavanagh *et al.* (2010) demonstrated that cold populations had a faster developmental rate and higher muscle mass than warm populations but at the cost of decreased development of skeletal structures. This finding may suggest that there are energetic costs associated with expressing different developmental rates but further studies are needed to evaluate whether such costs exist at the transcriptome level. Finally, plasticity can also be maladaptive and thus select against or compensate for faster evolutionary rate, a phenomenon known as counter gradient variation (Conover and Schultz 1995; Ghalambor et al. 2007).

### Modular gene expression response

The basic assumption underlying gene co-expression analyses is that functionally similar genes are likely co-expressed or their expression is correlated (Langfelder and Horvath 2008). We found evidence for the above assumption in the grayling transcriptome response to thermal treatments. We observed several enrichments for developmental traits among the modules, indicating that this approach can capture gene expression patterns underlying ecologically important traits. Previous studies have shown that graylings from cold-origin populations grow faster and have higher muscle mass than the warm origin populations (Kavanagh et al. 2010). We found muscle development related enrichments in the black module, which could be linked to the observed differentiation in the muscle growth patterns between cold and warm environments. In addition, we found several embryonic organ development enrichments in the turquoise module and they could be associated with several other early developmental traits that have differentiated in grayling populations (Haugen 2000; Haugen and Vøllestad 2000; Haugen and Vøllestad 2001; Koskinen et al. 2002). Similar results have been observed in lake whitefish (*Coregonus clupeaformis*), for which key phenotypic traits of adaptive significance were associated with co-expression modules (Filteau et al. 2013). Filteau *et al.* (2013) correlated phenotypic measurements to module eigengene expression, facilitating the direct association of ecologically important traits with gene expression patterns. We used a “top-down” approach, which can also enable the association of gene expression modules with previously identified adaptive traits in the grayling populations. In addition to enrichments associated with biological processes, we observed Pfam protein domain enrichment for Homeobox domain in the red module. The Hox gene cluster is a known transcription factor regulating embryonic development in the anterior posterior axis (e.g., Cheatle Jarvela & Hinman 2015). Originally discovered in Drosophila, Hox genes have been observed to control developmental processes in a wide variety of organisms (Pearson et al. 2005; Cheatle Jarvela and Hinman 2015). Recently, the Hox gene cluster was identified as a potential driver of diversification in coral reef fishes (Puebla et al. 2014). Hox genes control downstream genes through the up-regulation or down-regulation of gene expression and thus play an important role in regulating gene regulatory networks. However, additional studies are needed to elucidate the molecular mechanisms of how Hox genes control plastic or evolutionary responses in gene expression.

The modular pattern of gene expression suggests a flexible model of plastic and evolutionary responses during the early stages of thermal adaptation. Thus, the gene co-expression modules had variable responses to thermal treatment, population effect or their interaction or no effect at all. In general, plastic effects explained larger proportion of the eigengene expression than population effects. Overall, these results suggest that transcriptome is divided into subunits with separate biological functions and different evolutionary properties or gene expression responses. Although modularity is a characteristic of most living organisms at both the phenotypic and molecular levels, there is no consensus on the origin and maintenance of modularity (Espinosa-Soto and Wagner 2010). Several scenarios have been proposed based on computer simulations and empirical findings to explain how modularity evolves (Wagner 1996; Wagner et al. 2007; Espinosa-Soto and Wagner 2010). In general, modularity is expected to decrease pleiotropic interactions among the modules, thereby enabling more independent evolution of separate traits (Wagner et al. 2007; Snell-Rood et al. 2010). During adaptive evolution, most of the traits are under stabilizing selection, whereas a few traits are under directional selection (Wagner and Altenberg 1996; Espinosa-Soto and Wagner 2010). There is little evidence for the selection scenario in the present study. For example, the black module was enriched for muscle development genes, whereas the blue module was enriched for nervous system development genes, but both modules evolved under neutrality. Kavanagh *et al.* (2010) observed that the development of the musculoskeletal traits in grayling involved trade-offs. Faster muscle growth in the cold treatment likely constrains development of other traits, suggesting that genetic correlations might constrain the independent evolution of developmental modules. According to modularly varying evolutionary goals scenario, the modularly variable environment may trigger modular evolution. This idea has been demonstrated with computer simulations and in experimental studies (Parter et al. 2007; Espinosa-Soto and Wagner 2010). Parter *et al.* (2007) showed that bacterial populations living in variable environments showed more modular organization of metabolic networks than populations in stable environments. This scenario is appealing for examination in the grayling system to reveal habitat-specific modular patterns, but our attempts to construct robust population-specific gene co-expression modules resulted in low reproducibility of the modules. Espinosa-Soto & Wagner (2010) used computer simulations to show that modularity could arise as a by-product of selection favouring new gene activity patterns in certain organismal structures or under novel environmental conditions. Empirical findings support this scenario because most of the new evolutionary innovations are built on previously evolved modules (Espinosa-Soto and Wagner 2010). Demonstrating whether new gene activity patterns are underlying modular transcriptome evolution would require comparative data from several species (Espinosa-Soto and Wagner 2010). Finally, computer simulations indicate that maximizing network performance and minimizing connections costs can facilitate network modularity (Clune et al. 2013). In conclusion, the mechanism driving the origin and maintenance of the modularity in the present study cannot be inferred with certainty. However, we suggest that modularity may facilitate the flexible adjustment of gene expression levels to local thermal conditions as indicated module-specific plastic and evolutionary responses.

### No evidence for adaptive evolution in gene expression?

We detected no clear signals of adaptive evolution, suggesting that neutral patterns can explain gene expression variance after a recent colonization of varying thermal environments. We found only two transcripts outside the FST distribution (assuming c/h^2^=1.5 or 1), and remaining QST estimates fell within the F_ST_ distribution. These two transcripts were annotated to genes SSUH2 and tctex1d1. SSHU2 is involved in human dental malformations (Xiong et al. 2017), and tctex1d1 is associated with relative testis weight and is located in a genomic region of high sequence differentiation between house mouse (*Mus musculus*) subspecies (Phifer-Rixey et al. 2014). The overall pattern of gene expression divergence is consistent with neutral theory of evolution, predicting that genetic drift is expected to overdrive natural selection in small populations (Nei et al. 2010). Previous gene expression evolution studies have suggested that stabilizing selection and neutral evolution explains gene expression divergence between closely related species (Rifkin et al. 2003). For example, using a quantitative genetic model, Lemos *et al.* (2005) showed that 61-100% of the expressed genes in Drosophila species and mouse strains were under stabilizing selection, but little expression variance was explained by genetic drift or directional selection. Khaitovich (*et al.* 2004) examined primate gene expression, showing that expression differences accumulated linearly with time, consistent with neutral expectations. Previous studies using Q_ST_-F_ST_ comparisons have provided evidence for natural selection in gene expression data, but the majority of the gene expression divergence is consistent with neutral evolution (Roberge *et al.* 2007; Kohn *et al.* 2008; Aykanat *et al.* 2011; Papakostas *et al.* 2014; Leder *et al.* 2015). However, direct comparisons of Q_ST_-F_ST_ studies are not without problems (Leinonen et al. 2008; Leinonen et al. 2013). Previous studies have compared the mean and 95% confidence intervals of F_ST_ estimated from a few microsatellite markers to the Q_ST_ distribution, but this approach might be biased because the genome wide variance of F_ST_ could be underestimated (Whitlock 2008; Leinonen et al. 2013). Whitlock (2008) simulated F_ST_ and Q_ST_ distributions and observed that Q_ST_ and single locus F_ST_ distributions behave similarly under the Lewontin-Kraukauer model, suggesting that the entire distribution range of locus-specific F_ST_ estimates more realistically describes the neutral distribution. We used more than two thousand SNP markers, resulting in a more genome-wide perspective on variance of F_ST_. However, in our study, both estimators may suffer from sampling bias, reflecting small sample size and a low number of populations (O’Hara and Merilä 2005; Whitlock 2008), potentially resulting in a large sampling variance of both estimators. Theoretical studies have shown that a large number of samples and populations are needed to accurately estimate Q_ST_ and F_ST_ (O’Hara and Merilä 2005; Whitlock 2008). Furthermore, the heritability of gene expression considerably varies from gene to gene (Leder et al. 2015). The common garden design did not allow the estimation heritability, but the assumed ratios of additive variance and heritability indicated that Q_ST_-F_ST_ overlapped in a wide parameter range.

Similarly, direct comparisons of transcriptome divergence to previous studies demonstrating adaptive evolution in early-life history traits at the phenotypic level in grayling populations are slightly challenging. Koskinen (*et al.* 2002) and Kavanagh (*et al.* 2010) reported higher divergence than would be expected under neutrality in early life history traits, such as muscle growth, although in a different set of populations from the same region. Neutral divergence in gene expression was evident when the divergence in transcripts with homology to zebrafish muscle proteins and embryonic organ development were considered. The mean Q_ST_ was 0.041 for eleven muscle growth-related transcripts and 0.146 for embryonic organ development-related transcripts, indicating divergence consistent with neutrality. In addition to the statistical difficulties in estimating adaptive evolution in gene expression, further complications estimating and interpreting gene expression divergence compared to the phenotypic level might arise from the complexity of the molecular mechanisms underlying genotype-phenotype maps (Diz et al. 2012; Harrison et al. 2012; Alvarez et al. 2014). First, gene expression variability is inherently noisy because of environmental effects or effects arising from the maternal or paternal environment. Common garden experiments should in theory remove environmental effects, but trans generational plasticity (TGP) may bias estimating evolutionary responses in gene expression, even in the common environment (Salinas and Munch 2012; Shama et al. 2016). For example, Shama *et al.* (2016) showed that gene expression patterns in sticklebacks (*Gasterosteus aculeatus*) follow the environment experienced by the maternal environment, and these effects can persist for several generations. Salinas & Munch (2012) demonstrated that the parental rearing temperature modified the growth reaction norms in sheep head minnow offspring (*Cyprinodon variegatus*). Therefore, TGP could bias heritability estimates and lead to false conclusions about the rate of rapid adaptive evolution (Salinas and Munch 2012). Second, there is uncertainty about the gene expression variance that is functionally important or having a phenotypic effect, particularly when only quantitative data are available, as in many RNA-seq studies (Harrison et al. 2012). In Q_ST_-F_ST_ comparisons, the extreme values in the tails of the distribution are most likely candidates affected by natural selection, but variation falling within the neutral distribution might also have adaptive significance. Documented gene expression changes underlying adaptive traits can be relatively small, as in human hair colour variation (Guenther et al. 2014), or can involve almost complete tissue-specific silencing of expression, as in the pelvic spine loss in sticklebacks (Chan et al. 2010). Therefore, the Q_ST_-F_ST_ approach to analysing gene expression data to identify extreme values might not always be completely warranted. Finally, gene expression patterns depend on the topological features of a given network and the position of a gene in the network (Siegal et al. 2007; Levy and Siegal 2008). The network can buffer against the expression changes of individual genes to a certain degree, but highly connected genes or internal hub genes can be more vital to the entire network function and output (Han et al. 2004; Levy and Siegal 2008; Garfield et al. 2013). For example, the knockout of hub genes can be almost lethal in yeast (Han et al. 2004), and the number of protein interactions can constrain expression patterns. Therefore, estimating gene expression divergence should also consider the position of a gene in a network and the number of interactions with neighbouring genes.

## Conclusions

Our study revealed that each gene co-expression module varied in plastic and population responses. Overall, plastic responses explained a larger amount of the eigengene expression variance, suggesting that plasticity might be a key mechanism in adaptation to the local thermal conditions among these grayling populations. Plasticity showed population-specific responses, suggesting that plasticity might evolve according to patterns consistent with the Baldwin effect rather than genetic assimilation effect, where loss of plasticity is expected. Although populations showed signals of differentiation in expression profiles, no clear signals of adaptive evolution in gene expression were observed. The population differences were explained by patterns consistent with genetic drift, but sampling variance in both F_ST_ and Q_ST_ estimators because of low sample sizes might lead to low power of detecting selection. The modular organization of the gene expression patterns might enable module-specific tuning of gene expression to local thermal conditions. Overall, we suggest that combining systems and quantitative genetics methods can help in understanding the evolution of complex gene expression networks.

## Material and Methods

### Sample collection and common garden experiment

The study system comprises the ancestral river population and three derived small mountain lake populations located in central Norway (figure 1). The colonization history of these populations was inferred from historical records (Haugen and Vøllestad 2000; Haugen and Vøllestad 2001). The initial colonization of the L. Lesjaskogsvatnet occurred in the 1880s, when a temporary channel from the R. Gudbrandsdalslågen was opened. Therefore, the R. Gudbrandsdalslågen represents the ancestral grayling population of the system. From L. Lesjaskogsvatnet, a few grayling individuals were transported to high-elevation mountain lakes (L. Hårrtjønn and L. Øvre Merrabotvatnet) in the 1910s. A natural colonization from L. Hårrtjønn to L. Aursjøen occurred during the 1920s (Haugen and Vøllestad 2001) (figure 1). Thus, the divergence in this system occurred in the past 25-30 generations, assuming that the generation time for grayling is approximately six years (Haugen and Vøllestad 2001). The four study populations can be roughly classified as “warm” or “cold” origin populations according to the mean temperature during the grayling spawning season and early development period in June-August (Haugen 2000, supplementary figure 1, supplementary figure 2). In this respect, R. Gudbrandsdalslågen and L. Hårrtjønn can be described as “warm” origin populations, whereas L. Lesjaskogsvatnet and L. Aursjøen can be described as “cold” origin populations. The mean temperature differences translate to large temperature sums differences during the period of June-August (supplementary figure 1, supplementary figure 2).

The sample from R. Gudbrandsdalslågen was collected close to the town Otta, representing the ancestral population of the study system. The Otta location is below a present-day natural migration barrier to grayling, indicating that this population has probably been isolated from the other populations for hundreds of years (Junge et al. 2014). The L. Lesjaskogsvatnet sample was collected from R. Valåe, which is a small cold tributary in the eastern part of the lake. The sample from L. Hårrtjønn was collected from a small river outlet, and the sample from L. Aursjøen was collected from the main tributary (R. Kvita). Mature male and female fish were collected from each location during the 2013-spawning season for a semi common garden experiment. Eggs and sperm were extracted from mature fish under anaesthesia and subsequently transported on ice to the University of Oslo experimental facility. The experimental crosses were performed according to Thomassen *et al.* (2011). Briefly, for each population, eggs from 4-5 females were pooled and fertilized with sperm collected from 4-6 males. Individual fertilized eggs were subsequently transferred to standard 24-well culture plates with temperature-acclimated water added to the wells. The culture plates were incubated in climate-controlled rooms and at target temperatures of 6 °C and 10 °C. This design was thus a reciprocal thermal treatment, as these temperatures represent the average temperatures experienced by developing embryos in their natal environments (cold populations in 6 °C and warm populations in 10 °C). The number of degree days was used as a proxy for developmental stage to sample embryos from a same developmental stage (Chezik et al. 2014). Embryos were collected at ∼140 degree-days post fertilization, immediately individually frozen on dry ice in Eppendorf tubes, and subsequently stored at −80 °C until further analysis. Altogether 19 embryos from the cold treatment and 16 embryos from the warm treatment were sampled for the subsequent RNA-sequencing. The final sample set comprised five cold and four warm treatment embryos for Gudbrandsdalslågen, L. Hårrtjønn, and L. Aursjøen and four cold and four treatment embryos for L. Lesjagskogsvatnet.

### RNA extraction

RNA was extracted from whole embryos using TRI reagent according to the manufacturer’s instructions (Sigma-Aldrich). Before extraction, the tissue was homogenized using TissueLyser (Qiagen) for 30 s with full speed. The quality and quantity of the RNA were determined using a BioAnalyzer instrument (Agilent Technologies), and only samples with RNA integrity number (RIN) higher than eight were included in the sequencing. The sequencing libraries were prepared according to manufacturer’s instructions (Illumina). The sequencing was conducted at Beijing Genome Institute (BGI) using Illumina HiSeq 2000 equipment with 100-bp paired-end reads. To avoid lane effects, each sequencing library was distributed among five different lanes, and the reads were combined for subsequent analyses.

### Bioinformatic analyses

The quality of each sequencing library was investigated using the FastQC (v. 0.11.2) quality control tool for sequencing data (Andrews 2010). Analysis of the raw reads indicated the presence of low-quality bases in the 3’ end of the reads and an excess of PCR duplicates. ConDeTri read trimmer with default parameters was used to remove low-quality bases and PCR duplicates (Smeds and Künstner 2011). A *de novo* transcriptome assembly was reconstructed using all nine sequencing libraries (altogether c. 610 million reads) from the ancestral population R. Gudbrandsdalslågen. Before assembly, *in silico* normalization was used to restrict the maximum kmer coverage to 50x to decrease computational demands by reducing redundancy in the high-coverage regions. After normalization, 66.8 million reads remained for the *de novo* assembly. The *de novo* assembly was performed using the Trinity 2.0.4 assembler (Haas et al. 2013) with default parameters, except “minimum kmer coverage” was set to 10 and the “minimum glue” to 10. These parameters were adjusted to reduce the number of falsely identified transcripts as a result of low coverage, but the sensitivity for identifying lowly expressed transcripts can be lower compared to default parameters. The resulting transcripts were translated to proteins, and candidate-coding regions (or ORFs) were identified using TransDecoder software (http://transdecoder.github.io) with a minimum protein length of 100 amino acids. Similar protein sequences were merged using CD-HIT software (Li and Godzik 2006; Fu et al. 2012) with the sequence identity threshold set to one. The TransDecoder translated proteins (i.e., containing *in silico* predicted ORFs) were annotated using reciprocal protein-protein blast search using P-value cut-off 10^-5^ as implemented in BLAST+ software (Camacho et al. 2009). The blast searches were conducted against zebrafish (*Danio rerio*), stickleback (*Gasterosteus aculeatus*), cod (*Gadus morhua*) and Atlantic salmon (*Salmo salar*) protein databases. A local database for each species was constructed using protein annotations available in the Ensembl protein database (Cunningham et al. 2015) for zebrafish, stickleback and cod, and from GenBank (Benson et al. 2013) for Atlantic salmon. A transcript was considered reliably annotated when a significant reciprocal blast hit to one of the annotated proteins in any of the species was obtained. To compile a gene expression estimate or count table for each transcript, the quality filtered reads were mapped back to the *de novo* assembly. The mapping back was performed using Bowtie2 alignment software (Langmead and Salzberg 2012) with parameters -a -X 600 -rdg 6,5 -rfg 6,5 -score-min L, -0.6, -0.4 -no-discordant, -no-mixed. These parameters avoid mappings to splice variants and restrict the output to only read pairs mapped concordantly according to the eXpress software manual (http://bio.math.berkeley.edu/eXpress/faq.html). Transcript abundances, i.e., read counts per transcript, were estimated from the alignments using the default parameters in eXpress (Roberts and Pachter 2012). Rounded effective counts were used for the gene expression analyses as suggested in the eXpress manual. Effective counts are the expected number of reads generated from a given target (transcript), considering target length and the number of reads generated in the sequencing experiment (Roberts and Pachter 2012).

The Bowtie2 alignments described above were used for the identification of single nucleotide polymorphisms (SNPs). The *mpileup* command in SAMtools 1.4 package (Li et al. 2009) was used with a minimum mapping quality of 20 to combine mapping positions into a single file. SNPs and genotypes were called from the resulting pile up file using BCFtools and options –bcvg. The SNPs within 20 bp of indels and exceeding 2000x coverage were removed. Genotypes were filtered using minimum overall genotype quality 999, minimum overall genotype coverage 50, minimum individual genotype coverage 10, minimum number of samples of valid genotypes 35 (i.e. no missing data allowed) and overall minor allele frequency 5%. Loci deviating from Hardy-Weinberg equilibrium (both heterozygote excess and deficiency) at P-level 0.05 were removed. Finally, only one SNP per transcript was subjected to further analyses. Genetic differentiation was estimated using the Weir & Cockerham (1984) estimator of *F*_*ST*_ as implemented in the R package adegenet (Jombart and Ahmed 2011).

### Gene expression analyses

Potential unwanted variation in the count data arising from library preparation or other technical factors were investigated using principal component analysis. The R stats function *prcomp* in R statistical programming environment was used for PCA analysis. Three clusters were identified that explained 42.3% (PC1) and 14.1% (PC2) of the total variation and were not linked to the experimental setup, indicating the presence of possible unwanted variation (figure 2). To remove the unwanted variation in the data, residuals from a general linear model on non-normalized counts were used, and population and treatment were used as covariates (Risso et al. 2014). Briefly, this method should work for data without control genes, assuming that the covariates of interest are not correlated with unwanted variation (Risso et al. 2014). We used function *RUVr*, as implemented in the R package RUVSeq, to remove such effects (Risso et al. 2014). After removal of the unwanted variation, the samples were grouped according to population and treatment, and the PC1 and PC2 explained 25.9% and 17.3% of the total variation, respectively (figure 3).

A weighted gene co-expression analysis (WGCNA) was used to identify clusters of similarly expressed genes or modules using R package WGCNA (Langfelder and Horvath 2008). In this approach, similar gene co-expression patterns are identified based on expression correlation, which can subsequently be used to group transcripts into modules using hierarchical clustering (Langfelder and Horvath 2008). Gene expression variation within a module can be summarized to eigengenes using PCA, and variation in eigengenes can be linked to external information of interest (Langfelder and Horvath 2007; Langfelder and Horvath 2008). First outliers potentially interfering with network construction were detected using hierarchical clustering analysis with Euclidean distance to describe sample relationships. A sample from R. Gudbrandsdalslågen warm treatment was excluded due to a large distance from the other samples. A co-expression similarity matrix was calculated using signed (i.e., keeping the sign of co-expression) expression measures. The similarity matrix was transformed to adjacency matrix by raising the similarity between genes by to soft thresholding power of 13. This soft thresholding power was determined from the data, using a cut-off value of 0.9. In other words, genes were considered co-expressed if the correlation co-efficient exceeded 0.9 within a module. The other parameters used for the network construction were minimum module size 50 transcripts, deep split 2, and merge cut height 0.3. The stability of the modules was examined with 100 bootstrap replicates to assess the overlap of the module labels between non-sampled and re-sampled data sets. The overlap was estimated using Fisher’s exact test based on the module assignments. If the proportion of re-sampled data sets had significant overlap (p<0.01) more than 70%, then the module was considered statistically robust. The expression profile in each module was summarized to eigengene expression using the first principal component (PC1). The variation in eigengenes was analysed using ANOVA with population and treatment and their interaction as explanatory variables. The rationale for the ANOVA analyses is to detect plastic (treatment) and (population) responses. The ANOVA analyses were performed using the R stats function *aov*.

To estimate adaptive gene expression divergence, broad sense Q_ST_ (i.e., the additive genetic variance component is unknown) was estimated across the four study populations for all transcripts. In experimental settings where heritability or additive genetic variance cannot be estimated, Q_ST_ can be approximated assuming ratios of c/h^2^, where c represents the assumed proportion of total variance due to additive genetic effects among populations, and h^2^ represents heritability (Brommer 2011).

Transcriptome derived SNPs were used to estimate F_ST_ to obtain a neutral baseline to which Q_ST_ can be compared. The rationale for the Q_ST_-F_ST_ comparisons is to identify candidate transcripts under stabilizing or divergent selection (Leinonen et al. 2013). If a given transcript shows lower or higher differentiation compared to the F_ST_ distribution, then stabilizing and divergent selection can be inferred, respectively (Leinonen et al. 2013). Q_ST_ was calculated according to the formula 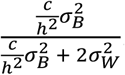, where 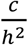 is the assumed ratio of additive variance and heritability, 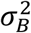 is the variance between populations in transcript expression and 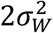 is the variance within population in transcript expression (Brommer 2011). The within and between population variance components in transcript expression were estimated fitting a mixed effect model using thermal treatment as a fixed effect and population as a random effect. The mixed effect model was fitted using *lme* function in R package lme4 (Bates et al. 2015). The confidence intervals of each Q_ST_ estimate were estimated with 250 bootstrap replicates. Three c/h^2^ ratios were assumed: 0.5, 1 and 1.5. The null assumption c/h^2^=1 assumes that the additive phenotypic variance between and within populations is the same, but this ratio can be smaller (0.5) or larger (1.5), reflecting, for example, environmental effects (Brommer 2011). The Q_ST_ estimates were compared to the entire distribution range of locus specific F_ST_ (proxy for the neutral distribution) according to Whitlock (2008). Transcripts showing a higher divergence (Q_ST_ > F_ST_) expected by genetic drift alone are potentially under directional selection whereas transcripts showing lower divergence (Q_ST_ < F_ST_) under balancing selection. Neutrally evolving transcripts are expected to fall within the F_ST_ distribution (Leinonen et al. 2013). The 95% confidence intervals of the Q_ST_ estimates were considered in the above comparisons. If the upper or lower confidence interval did not overlap with the lower or upper tail of F_ST_ distribution, then the given transcript was considered affected by stabilizing or divergent selection, respectively.

### Gene Ontology enrichment analyses

Gene enrichment analyses were performed using zebrafish gene annotations for each statistically robust module identified in the WGCNA analysis. The STRING database was used to identify significant (FDR < 0.05) Gene Ontology (GO) categories for biological processes and PFAM protein domains and features (Szklarczyk et al. 2015). The GO categories were summarized using SimRel semantic similarity measure to avoid interpretation of redundant categories. The merging of semantically similar GO categories was based on hierarchical clustering with a user-specified cutoff value C. A cutoff value 0.5 was used to merge similar categories, corresponding to 1% chance of merging two randomly generated categories (Supek et al. 2011). The p-values of the initial enrichment analyses were used to select a representative GO term for each merged category. Thus, the lowest p-value among the merged categories was selected as the representative GO term. The REVIGO web-server tool was used for semantic similarity analyses (Supek et al. 2011).

## Acknowledgements

The present study was financially supported through grants from the Finnish Academy (Project numbers 287342 and 302873) and the Norwegian Research Council (Project number 177728) and the Finnish Cultural Foundation. The authors thank Tutku Aykanat for assistance with QST-FST comparisons, and Ane Kvinge for assistance with field sampling. We acknowledge the CSC – IT Center for Science in Finland for providing computational resources.

---

**Supplementary figure 1.**
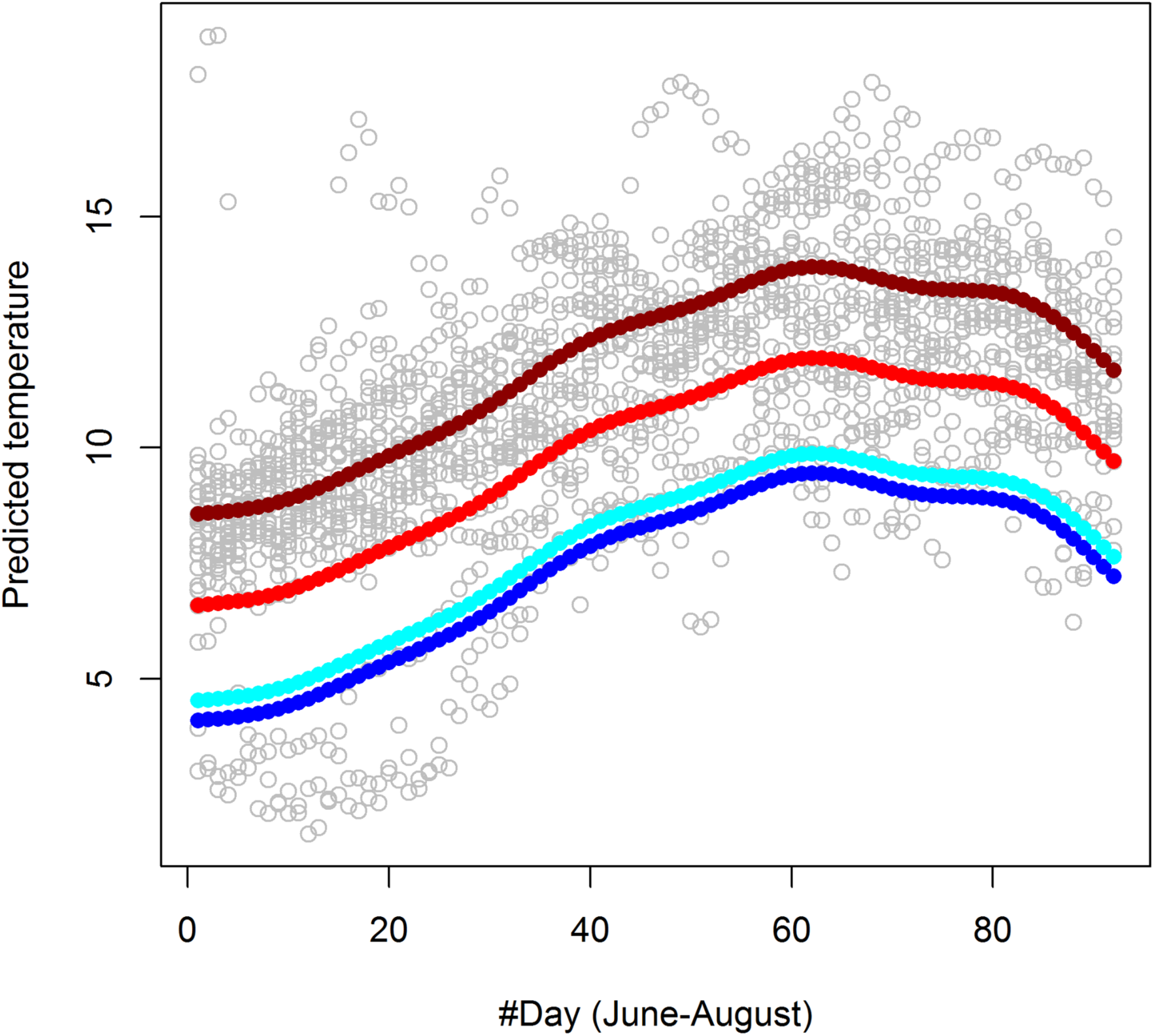
Temperature profiles of the four study populations during the grayling early developmental period during June-August. Generalized additive model predictions were used to estimate smoothed temperature changes in the study locations. Gam function in R package mgvc 1.8.17 was used to fit the model temperature ∼ location + s (day), where s is the covariate (the day rank since June 1st) used as a smoothing term. The data obtained from L. Hårrtjønn are from 1995 because of logistic difficulties in recovering the temperature logger in autumn 2013.

**Supplementary figure 2.**
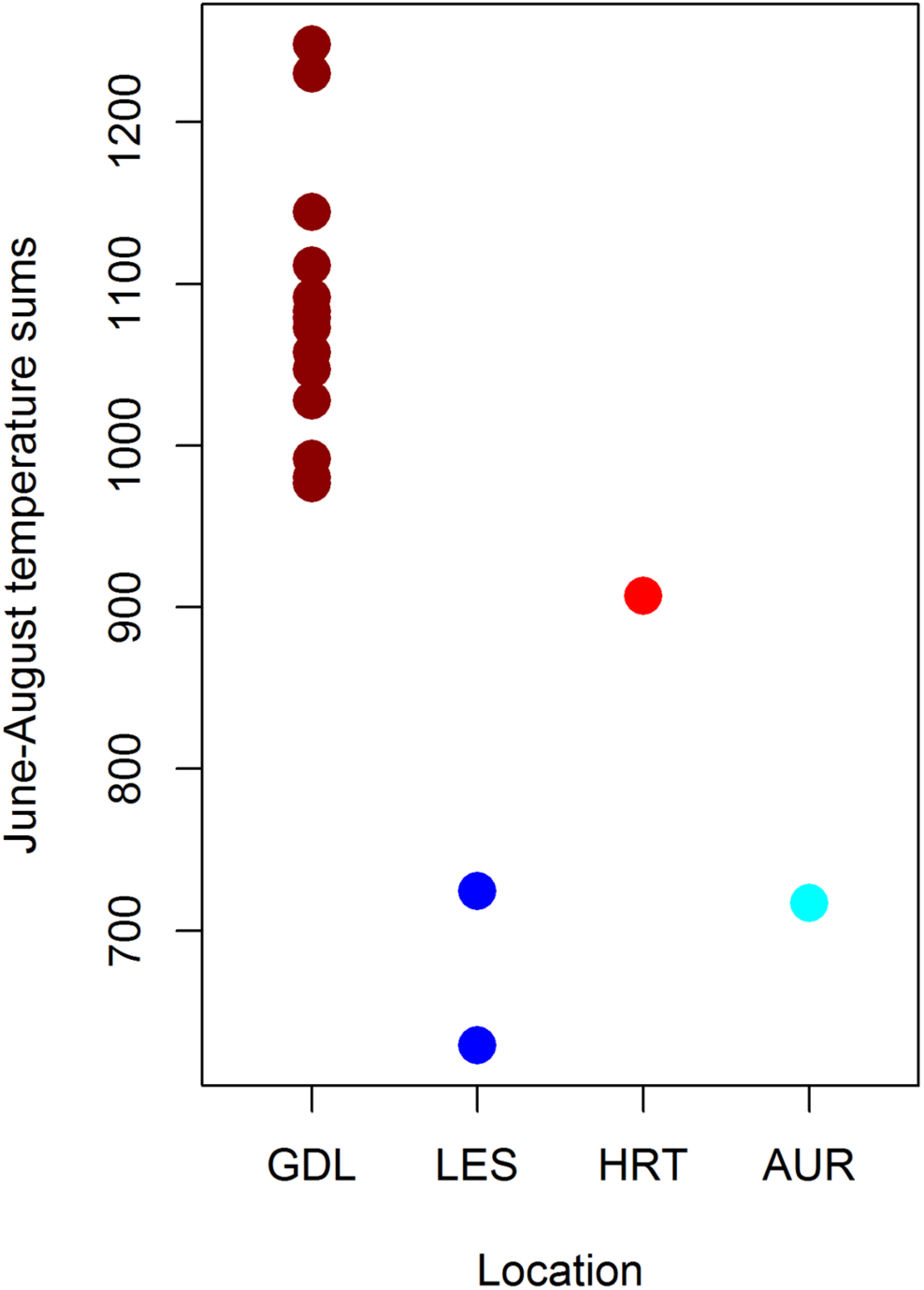
Temperature sums for June-August for each year and each location. Each temperature sum was calculated by summing the mean daily temperatures with max temperature. Missing data for two days (of the total 92) were imputed using the mean temperature. L. Hårrtjonn only had three measurements from June; thus, the predicted values from the gam model were used to fill in the missing data.

**Supplementary figure 3.**
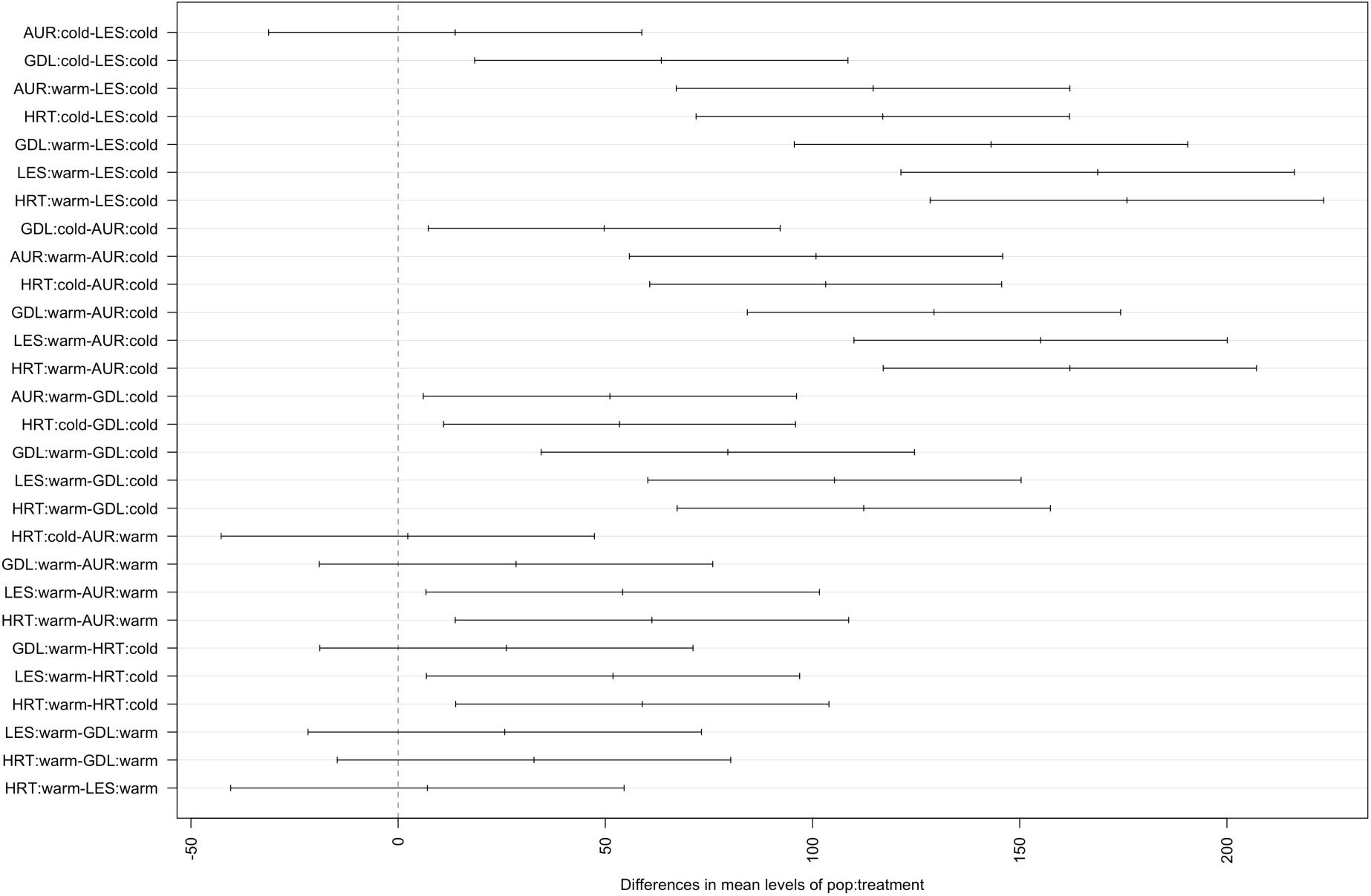
Tukey HSD plots showing the confidence intervals of the estimated effects of population and treatment on PC1.

**Supplementary figure 4.**
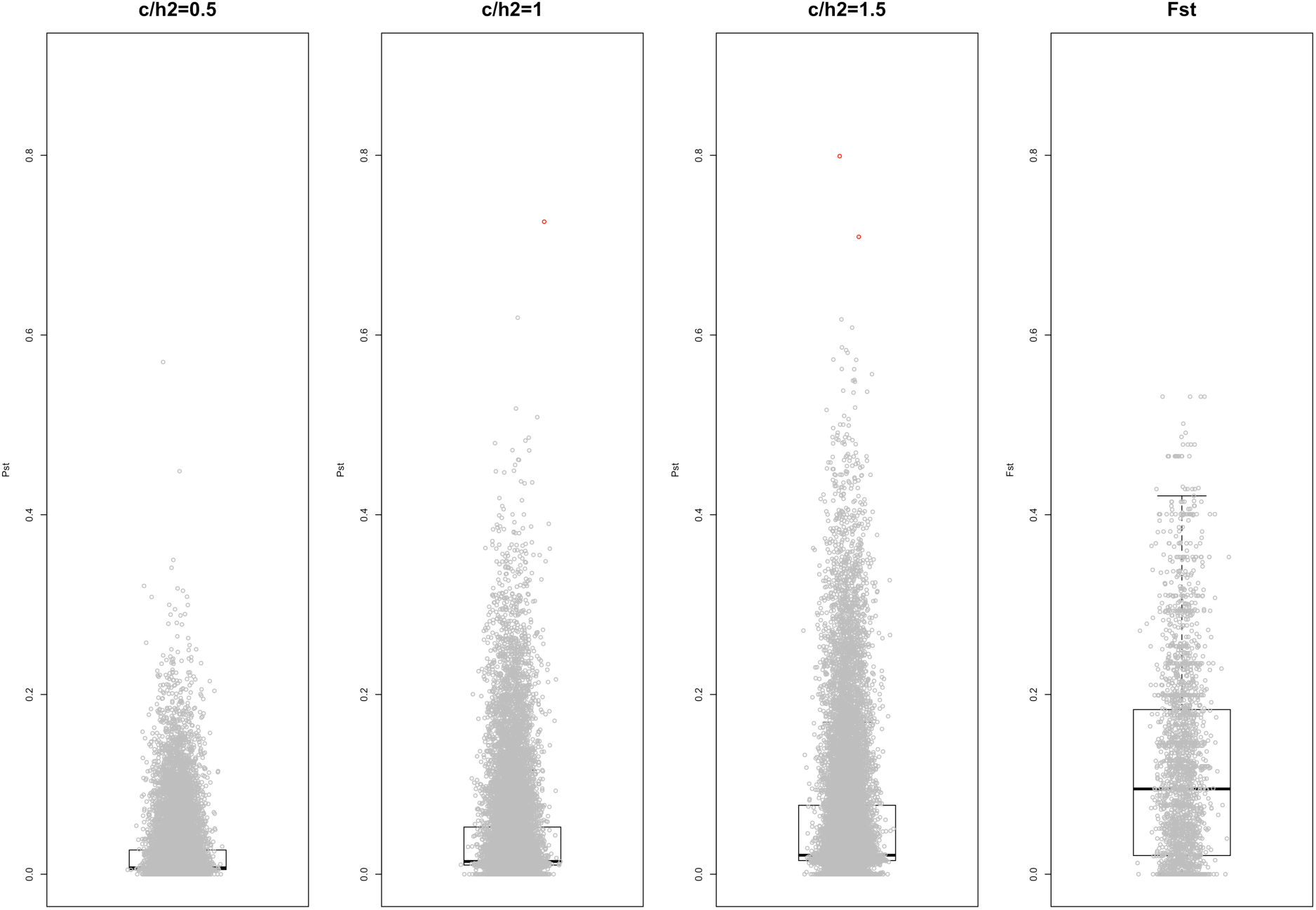
Q_ST_-F_ST_ comparisons showing overlap between the distributions across all transcripts. For Q_ST_, three scaling factors (c/h2), 0.5, 1 and 1.5, were assumed. Two transcripts outside the F_ST_ distribution are shown as red circles.

**Supplementary figure 5.**
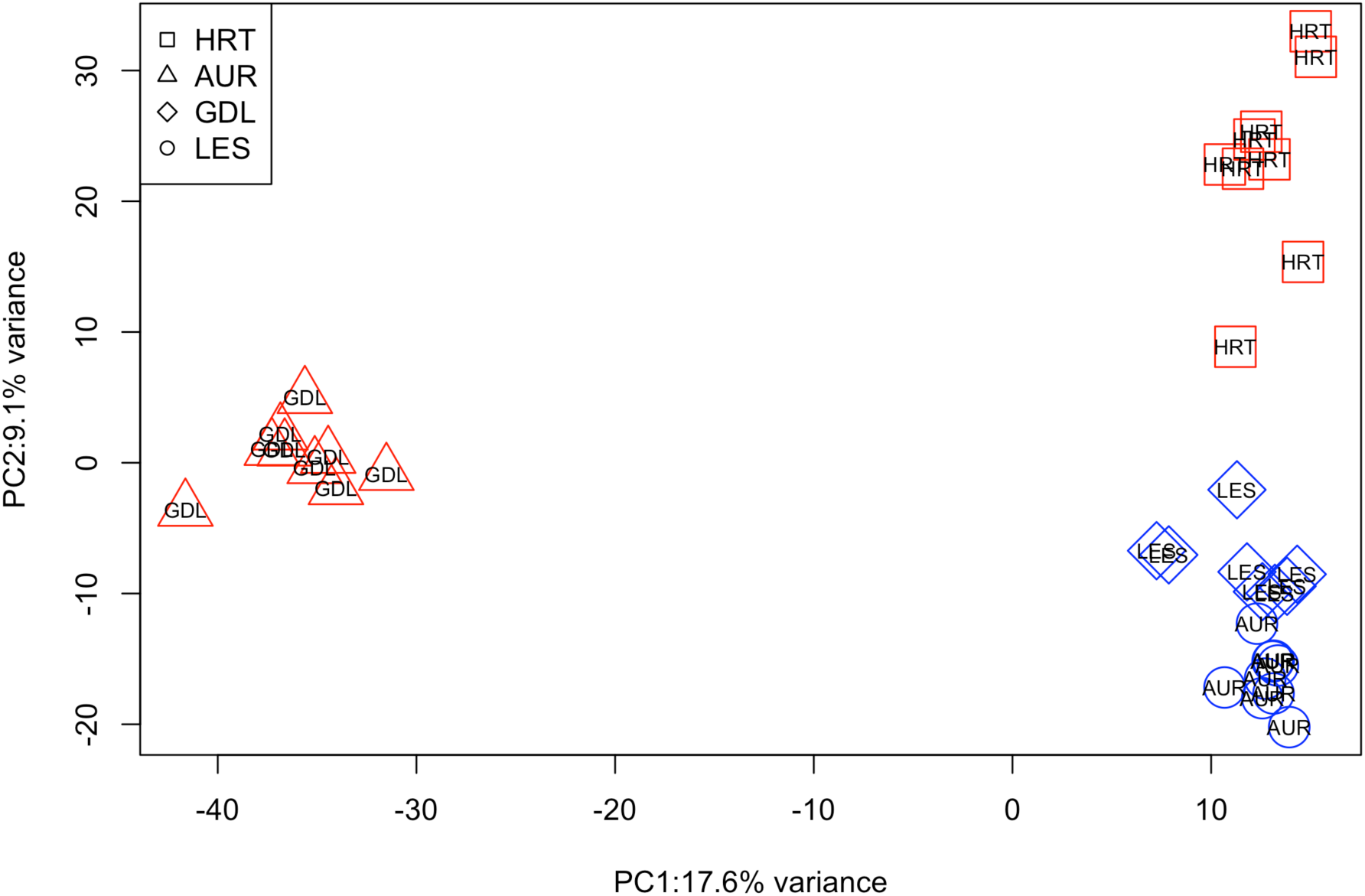
Genetic relationships (PCA) among the four study populations estimated from 2458 transcriptome-derived SNP loci. The colours indicate the thermal origin (blue = cold, red = warm).

**Supplementary figures 6-9.**
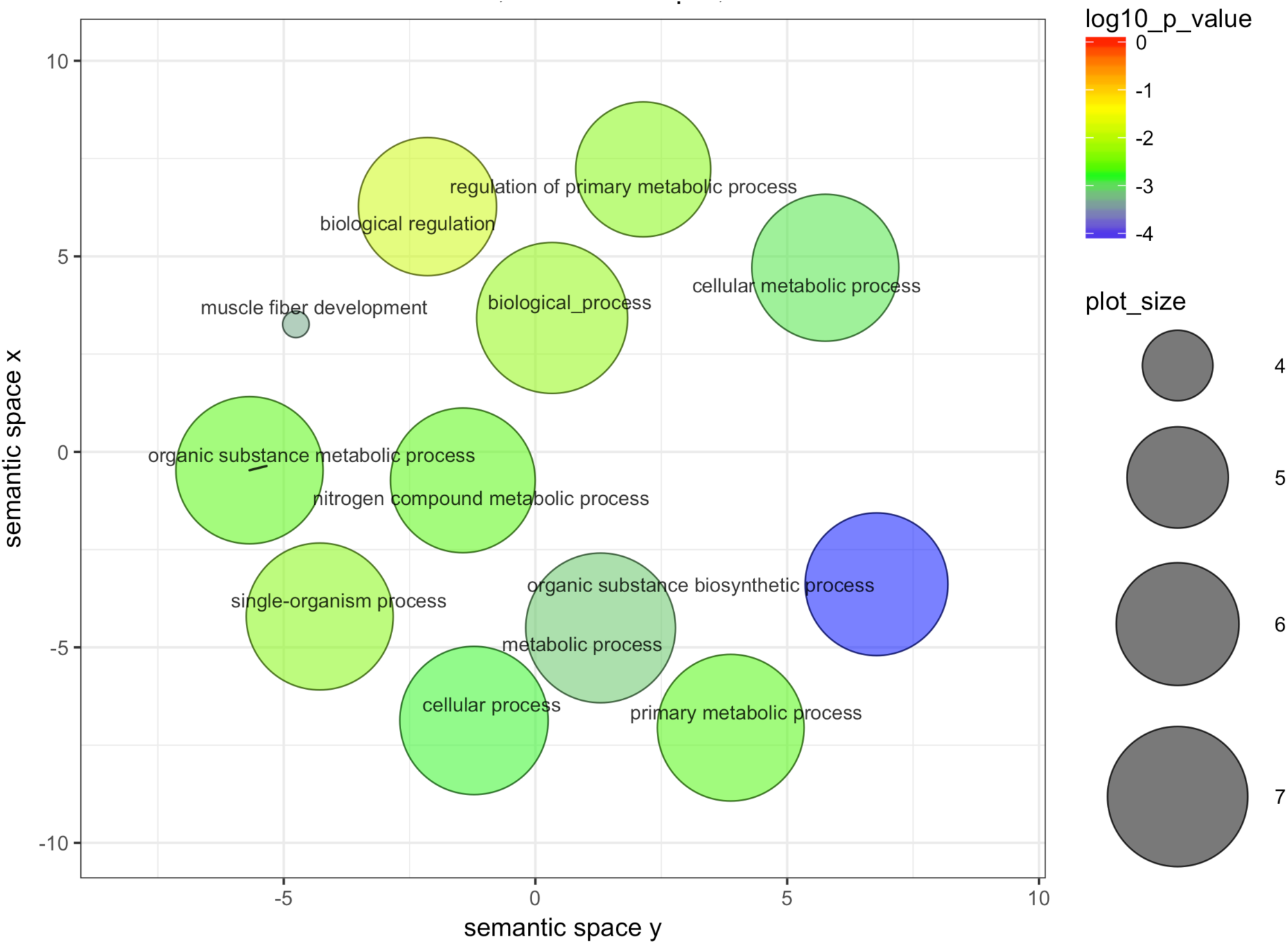
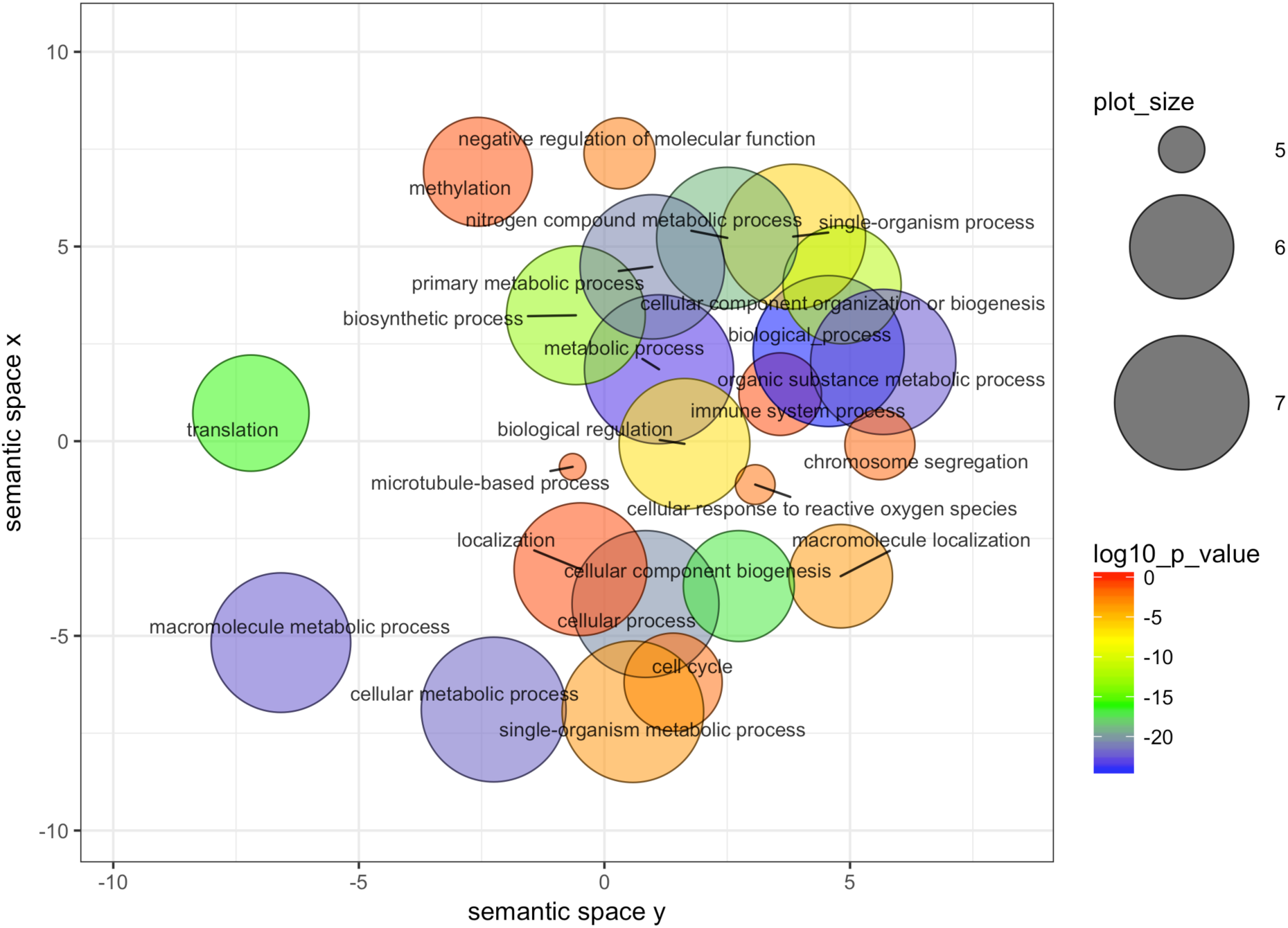
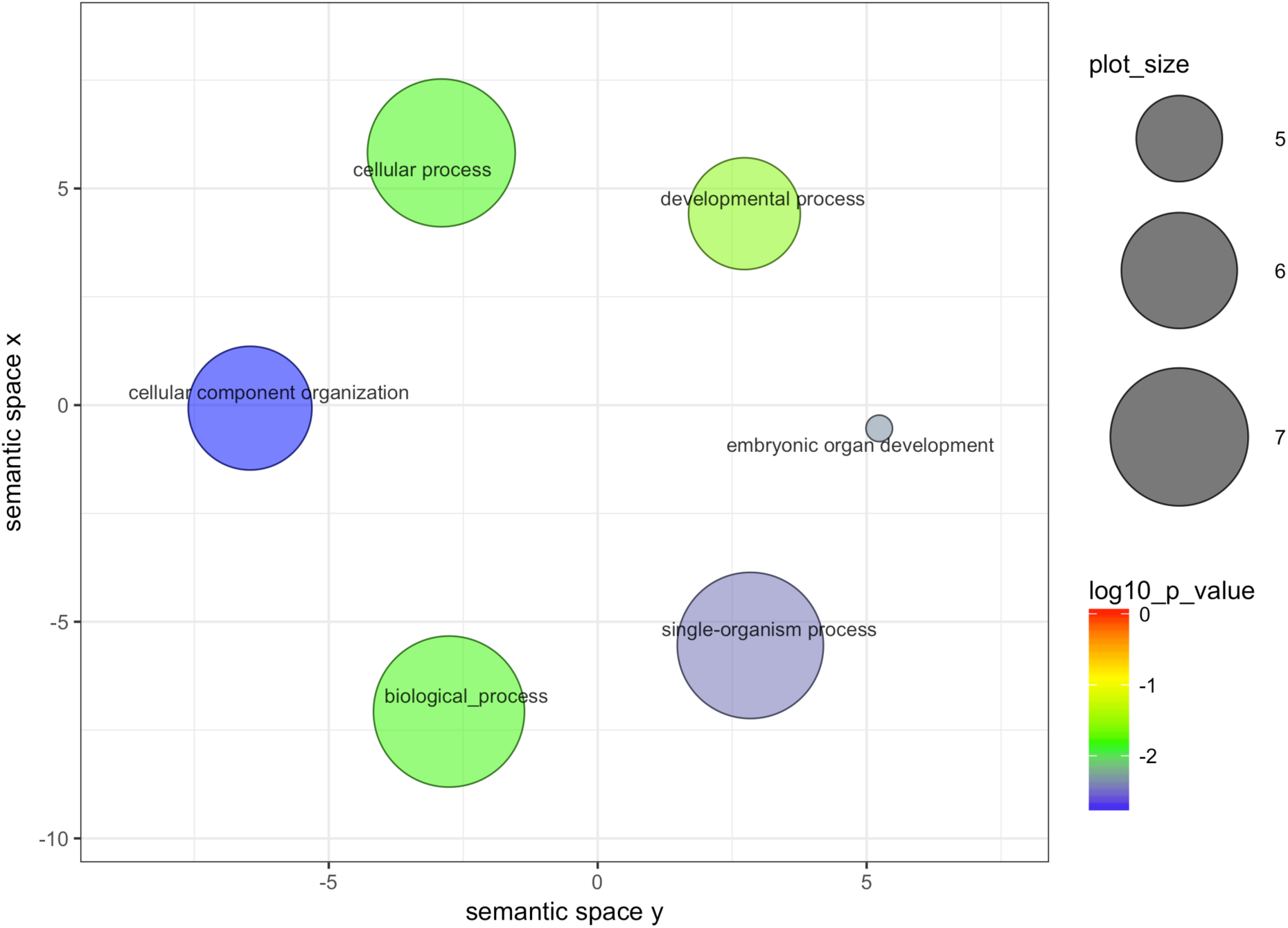
Gene enrichments after merging semantically similar GO categories. The number of transcripts assigned to each module and the number of annotated transcripts to zebrafish are shown in the headline. The size of the circle is proportional to the uniqueness of a given category in the Gene Ontology database. Smaller circles refer to more unique categories, whereas larger circles refer to broader categories.

**Supplementary table 1.** Summary statistics of the bioinformatic analyses and sampling origins and thermal treatments. QC filtered and QC% column indicate the number and percentage of remaining reads after removing low quality reads and PCR duplicates. Mapped reads and Mapping% columns show the number of reds mapped back to the de novo assembly.

| Library | Population | Treatment | Raw reads | QC filtered | QC% | Mapped reads | Mapping % |
| --- | --- | --- | --- | --- | --- | --- | --- |
| HA10-2 | HRT | 10 °C | 79769702 | 67636034 | 84.79 | 27252770 | 41.39 |
| HA10-3 | HRT | 10 °C | 78608284 | 67076326 | 85.33 | 27994128 | 41.73 |
| HA10-4 | HRT | 10 °C | 77698028 | 67902228 | 87.39 | 26655606 | 39.26 |
| HA10-5 | HRT | 10 °C | 78015312 | 65836598 | 84.39 | 27075006 | 39.18 |
| HA6-1 | HRT | 6 °C | 77596180 | 69104080 | 89.06 | 27337410 | 41.15 |
| HA6-2 | HRT | 6 °C | 76897538 | 66440580 | 86.40 | 26847766 | 41.3 |
| HA6-3 | HRT | 6 °C | 76939314 | 65004000 | 84.49 | 28999956 | 42.73 |
| HA6-4 | HRT | 6 °C | 79151454 | 67868918 | 85.75 | 26315166 | 39.71 |
| HA6-5 | HRT | 6 °C | 79677366 | 71323436 | 89.52 | 27541196 | 40.72 |
| KV10-2 | AUR | 10 °C | 78433640 | 66272138 | 84.49 | 26914788 | 40.3 |
| KV10-3 | AUR | 10 °C | 77515912 | 66789586 | 86.16 | 28347758 | 41.1 |
| KV10-4 | AUR | 10 °C | 80626390 | 68970452 | 85.54 | 26766552 | 39.57 |
| KV10-5 | AUR | 10 °C | 78081584 | 67646314 | 86.64 | 27434376 | 40.6 |
| KV6-1 | AUR | 6 °C | 77812126 | 67570588 | 86.84 | 26391102 | 38.37 |
| KV6-2 | AUR | 6 °C | 77392548 | 68775130 | 88.87 | 27287602 | 38.67 |
| KV6-3 | AUR | 6 °C | 79672922 | 70563186 | 88.57 | 28290174 | 39.25 |
| KV6-4 | AUR | 6 °C | 80833148 | 72085838 | 89.18 | 26770350 | 38.86 |
| KV6-5 | AUR | 6 °C | 79112344 | 68885708 | 87.07 | 28029008 | 40.39 |
| OT10-2 | GDL | 10 °C | 82035404 | 69389882 | 84.59 | 27177414 | 41.33 |
| OT10-3 | GDL | 10 °C | 78494188 | 65755420 | 83.77 | 27454262 | 38.9 |
| OT10-4 | GDL | 10 °C | 77541912 | 66327276 | 85.54 | 27194462 | 41.14 |
| OT10-5 | GDL | 10 °C | 80122586 | 67856496 | 84.69 | 27035380 | 38.51 |
| OT6-1 | GDL | 6 °C | 79588420 | 70584690 | 88.69 | 26633510 | 39.3 |
| OT6-2 | GDL | 6 °C | 77426760 | 65417540 | 84.49 | 27227732 | 39.82 |
| OT6-3 | GDL | 6 °C | 78405954 | 66097250 | 84.30 | 27131120 | 40.82 |
| OT6-4 | GDL | 6 °C | 77313850 | 69259894 | 89.58 | 27173410 | 39.14 |
| OT6-5 | GDL | 6 °C | 79481338 | 70203658 | 88.33 | 27498946 | 40.5 |
| VA10-1 | LES | 10 °C | 77385160 | 65443516 | 84.57 | 27082290 | 38.2 |
| VA10-2 | LES | 10 °C | 78840934 | 67906314 | 86.13 | 27539124 | 38.5 |
| VA10-3 | LES | 10 °C | 77038638 | 66990920 | 86.96 | 26496838 | 39.95 |
| VA10-4 | LES | 10 °C | 82600348 | 67622894 | 81.87 | 28567436 | 42.1 |
| VA6-1 | LES | 6 °C | 79693866 | 70557214 | 88.54 | 27358746 | 41.82 |
| VA6-2 | LES | 6 °C | 77579016 | 67143946 | 86.55 | 26575788 | 38.37 |
| VA6-3 | LES | 6 °C | 79651594 | 70887420 | 89.00 | 26312936 | 40.21 |
| VA6-5 | LES | 6 °C | 79935034 | 71525202 | 89.48 | 27619374 | 38.72 |
|  | Mean |  | 78770536 | 68134876 | 86.50 | 27307622 | 40.11 |

**Supplementary table 2.**
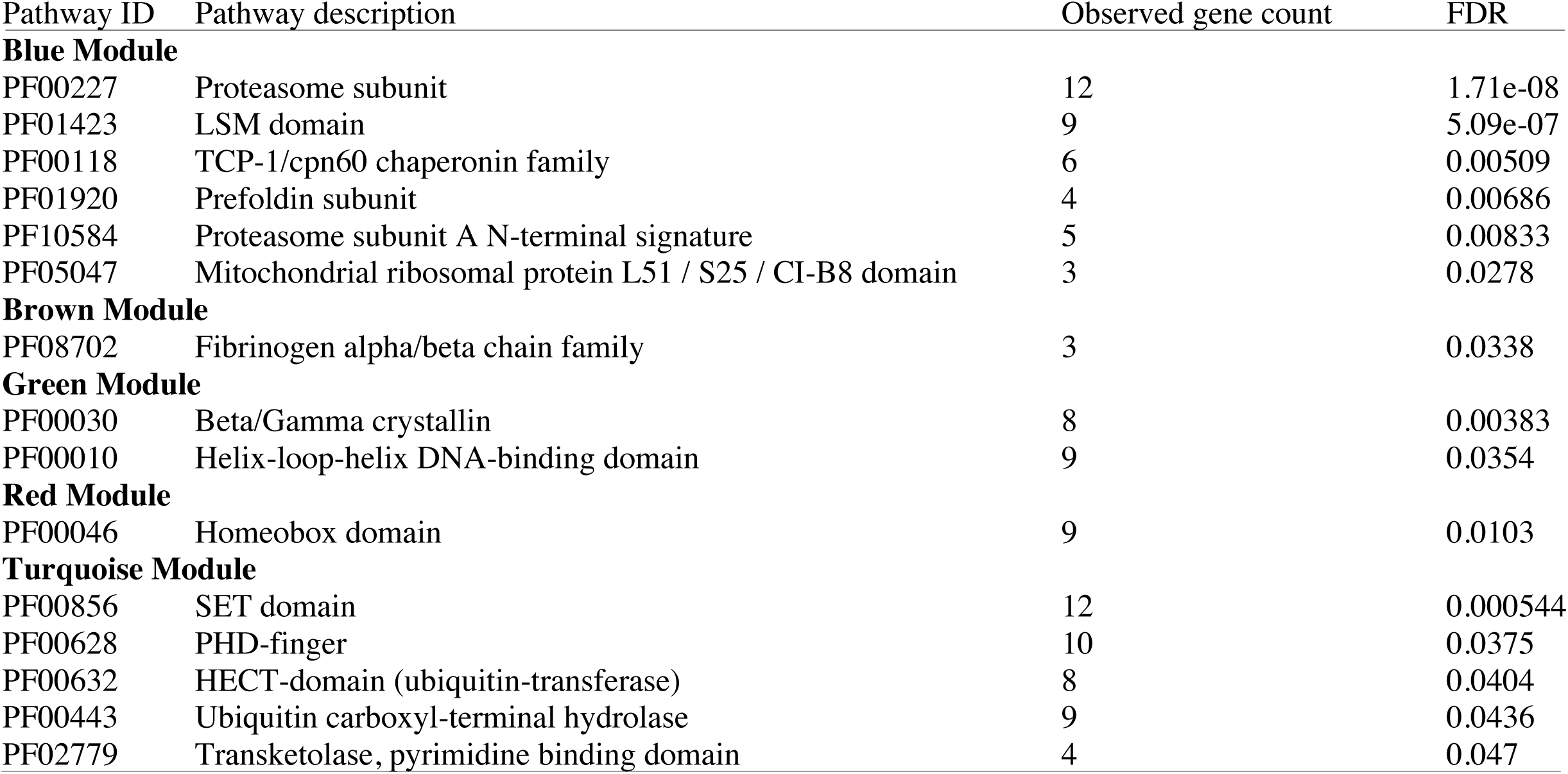
Pfam protein domain enrichments observed in the gene expression modules. The observed gene count indicates the number of zebrafish annotations in the given enrichments.

## References

Alvarez M, Schrey AW, Richards CL. 2014. Ten years of transcriptomics in wild populations: what have we learned about their ecology and evolution? Mol. Ecol. 24:710–725.

Aubin-Horth N, Renn SCP. 2009. Genomic reaction norms: Using integrative biology to understand molecular mechanisms of phenotypic plasticity. Mol. Ecol. 18:3763–3780.

Aykanat T, Thrower FP, Heath DD. 2011. Rapid evolution of osmoregulatory function by modification of gene transcription in steelhead trout. Genetica 139:233–242.

Bates D, Mächler M, Bolker B, Walker S. 2015. Fitting Linear Mixed-Effects Models using lme4. J. Stat. Softw. 67:51.

Benson DA, Cavanaugh M, Clark K, Karsch-Mizrachi I, Lipman DJ, Ostell J, Sayers EW. 2013. GenBank. Nucleic Acids Res. 41:36–42.

Brommer JE. 2011. Whither Pst? The approximation of Qst by Pst in evolutionary and conservation biology. J. Evol. Biol. 24:1160–1168.

Camacho C, Coulouris G, Avagyan V, Ma N, Papadopoulos J, Bealer K, Madden TL. 2009. BLAST plus: architecture and applications. BMC Bioinformatics 10:1.

Chan YF, Marks ME, Jones FC, Villarreal G, Shapiro MD, Brady SD, Southwick AM, Absher DM, Grimwood J, Schmutz J, et al. 2010. Adaptive evolution of pelvic reduction in sticklebacks by recurrent deletion of a Pitx1 enhancer. Science 327:302–305.

Cheatle Jarvela AM, Hinman VF. 2015. Evolution of transcription factor function as a mechanism for changing metazoan developmental gene regulatory networks. Evodevo 6:3.

Chevin LM, Lande R, Mace GM. 2010. Adaptation, plasticity, and extinction in a changing environment: Towards a predictive theory. PLoS Biol. 8.

Chezik KA., Lester NP, Venturelli PA. 2014. Fish growth and degree-days I: selecting a base temperature for a within-population study. Can. J. Fish. Aquat. Sci. 55:47–55.

Clune J, Mouret J-B, Lipson H. 2013. The evolutionary origins of modularity. Proc. Biol. Sci. 280:20122863.

Conover DO, Schultz ET. 1995. Phenotypic similarity and the evolutionary significance of countergradient variation. Trends Ecol. Evol. 10:248–252.

Crispo E. 2007. The Baldwin effect and genetic assimilation: Revisiting two mechanisms of evolutionary change mediated by phenotypic plasticity. Evolution. 61:2469–2479.

Crozier LG, Hutchings JA. 2014. Plastic and evolutionary responses to climate change in fish. Evol. Appl. 7:68–87.

Cunningham F, Amode MR, Barrell D, Beal K, Billis K, Brent S, Carvalho-Silva D, Clapham P, Coates G, Fitzgerald S, et al. 2015. Ensembl 2015. Nucleic Acids Res. 43:D662–D669.

DeBiasse MB, Kelly MW. 2016. Plastic and Evolved Responses to Global Change: What Can We Learn from Comparative Transcriptomics? J. Hered. 107:71–81.

DeWitt TJ, Sih A, Wilson DS. 1998. Costs and limits of phenotypic plasticity. Trends Ecol. Evol. 13:77–81.

Diz AP, Martínez-Fernández M, Rolán-Alvarez E. 2012. Proteomics in evolutionary ecology: linking the genotype with the phenotype. Mol. Ecol. 21:1060–1080.

Draghi JA, Whitlock MC. 2012. Phenotypic plasticity facilitates mutational variance, genetic variance, and evolvability along the major axis of environmental variation. Evolution. 66:2891–2902.

Ehrenreich IM, Pfennig DW. 2015. Genetic assimilation: a review of its potential proximate causes and evolutionary consequences. Ann. Bot 117:769–779.

Espinosa-Soto C, Wagner A. 2010. Specialization can drive the evolution of modularity. PLoS Comput. Biol. 6.

Feltus FA. 2014. Systems genetics: a paradigm to improve discovery of candidate genes and mechanisms underlying complex traits. Plant Sci. 223:45–48.

Fierst JL. 2011. A history of phenotypic plasticity accelerates adaptation to a new environment. J. Evol. Biol. 24:1992–2001.

Filteau M, Pavey SA, St-Cyr J, Bernatchez L. 2013. Gene coexpression networks reveal key drivers of phenotypic divergence in lake whitefish. Mol. Biol. Evol. 30:1384–1396.

Fischer EK, Ghalambor CK, Hoke KL. 2016. Can a Network Approach Resolve How Adaptive vs Nonadaptive Plasticity Impacts Evolutionary Trajectories. Integr. Comp. Biol. 56:877–888.

Forsman A. 2015. Rethinking phenotypic plasticity and its consequences for individuals, populations and species. Heredity 115:276–284.

Franks SJ, Hoffmann AA. 2012. Genetics of climate change adaptation. Annu. Rev. Genet. 46:185–208.

Fraser HB. 2011. Prospects & Overviews Genome-wide approaches to the study of adaptive gene expression evolution. BioEssays 33:469–477.

Friedrich SR, Meyer A. 2016. How plasticity, genetic assimilation and cryptic genetic variation may contribute to adaptive radiations. Mol. Ecol. 26:330–350.

Fu L, Niu B, Zhu Z, Wu S, Li W. 2012. CD-HIT: Accelerated for clustering the next-generation sequencing data. Bioinformatics 28:3150–3152.

Fusco G, Minelli A. 2010. Phenotypic plasticity in development and evolution: facts and concepts. Philos. Trans. R. Soc. London B Biol. Sci. 365:547:556.

Garfield DA, Runcie DE, Babbitt CC, Haygood R, Nielsen WJ, Wray GA. 2013. The Impact of Gene Expression Variation on the Robustness and Evolvability of a Developmental Gene Regulatory Network.Hurst LD, editor. PLoS Biol. 11:e1001696.

Ghalambor CK, Hoke KL, Ruell EW, Fischer EK, Reznick DN, Hughes K a. 2015. Non-adaptive plasticity potentiates rapid adaptive evolution of gene expression in nature. Nature 525:372–375.

Ghalambor CK, McKay JK, Carroll SP, Reznick DN. 2007. Adaptive versus non-adaptive phenotypic plasticity and the potential for contemporary adaptation in new environments. Funct. Ecol. 21:394–407.

Gienapp P, Teplitsky C, Alho JS, Mills JA, Merilä J. 2008. Climate change and evolution: disentangling environmental and genetic responses. Mol. Ecol. 17:167–178.

Gregersen F, Haugen TO, Vøllestad LA. 2008. Contemporary egg size divergence among sympatric grayling demes with common ancestors. Ecol. Freshw. Fish 17:110–118.

Guenther CA, Tasic B, Luo L, Bedell MA, Kingsley DM. 2014. A molecular basis for classic blond hair color in Europeans. Nat. Genet. 46:748–752.

Haas BJ, Papanicolaou A, Yassour M, Grabherr M, Blood PD, Bowden J, Couger MB, Eccles D, Li B, Lieber M, et al. 2013. De novo transcript sequence reconstruction from RNA-seq using the Trinity platform for reference generation and analysis. Nat. Protoc. 8:1494–1512.

Han J-DJ, Bertin N, Hao T, Goldberg DS, Berriz GF, Zhang L V., Dupuy D, Walhout AJM, Cusick ME, Roth FP, et al. 2004. Evidence for dynamically organized modularity in the yeast protein– protein interaction network. Nature 430:88–93.

Harrison PW, Wright AE, Mank JE. 2012. The evolution of gene expression and the transcriptome-phenotype relationship. Semin. Cell Dev. Biol. 23:222–229.

Haugen T, Vøllestad LA. 2000. Population differences in early life-history traits in grayling. J. Evol.Biol. 13:897–905.

Haugen TO. 2000. Growth and survival effects on maturation pattern in populations of grayling with recent common ancestors. J. Fish Biol. 56:1173–1191.

Haugen TO, Vøllestad LA. 2001. A century of life-history evolution in grayling. Genetica 112–113:475–491.

Hendry AP. 2016. Key Questions on the Role of Phenotypic Plasticity in Eco-Evolutionary Dynamics. J. Hered. 107:25–41.

Hoekstra HE, Coyne JA. 2007. The locus of evolution: Evo Devo and the genetics of adaptation. Evolution. 61:995–1016.

Jombart T, Ahmed I. 2011. adegenet 1.3-1: new tools for the analysis of genome-wide SNP data. Bioinformatics 27:3070–3071.

Junge C, Museth J, Hindar K, Kraabøl M, Vøllestad LA. 2014. Assessing the consequences of habitat fragmentation for two migratory salmonid fishes. Aquat. Conserv. Mar. Freshw. Ecosyst. 24:297–311.

Kavanagh KD, Haugen TO, Gregersen F, Jernvall J, Vøllestad LA. 2010. Contemporary temperature-driven divergence in a Nordic freshwater fish under conditions commonly thought to hinder adaptation. BMC Evol. Biol. 10:350.

Khaitovich P, Weiss G, Lachmann M, Hellmann I, Enard W, Muetzel B, Wirkner U, Ansorge W, Pääbo S. 2004. A neutral model of transcriptome evolution. PLoS Biol. 2:E132.

Kohn MH, Shapiro J, Wu C-I. 2008. Decoupled differentiation of gene expression and coding sequence among Drosophila populations. Genes Genet. Syst. 83:265–273.

Koskinen MT, Haugen TO, Primmer CR. 2002. Contemporary fisherian life-history evolution in small salmonid populations. Nature 419:826–830.

Laarits T, Bordalo P, Lemos B. 2016. Genes under weaker stabilizing selection increase network evolvability and rapid regulatory adaptation to an environmental shift. J. Evol. Biol. 29:1602–1616.

Lande R. 2015. Evolution of phenotypic plasticity in colonizing species. Mol. Ecol. 24:2038–2045.

Langfelder P, Horvath S. 2007. Eigengene networks for studying the relationships between co-expression modules. BMC Syst. Biol. 1:54.

Langfelder P, Horvath S. 2008. WGCNA: an R package for weighted correlation network analysis. BMC Bioinformatics 9:559.

Langmead B, Salzberg SL. 2012. Fast gapped-read alignment with Bowtie 2. Nat. Methods 9:357–359.

Leder EH, McCairns RJS, Leinonen T, Cano JM, Viitaniemi HM, Nikinmaa M, Primmer CR, Merilä J. 2015. The evolution and adaptive potential of transcriptional variation in sticklebacks-signatures of selection and widespread heritability. Mol. Biol. Evol. 32:674–689.

Leinonen T, McCairns RJS, O’Hara RB, Merilä J. 2013. Q(ST)-F(ST) comparisons: evolutionary and ecological insights from genomic heterogeneity. Nat. Rev. Genet. 14:179–190.

Leinonen T, O’Hara RB, Cano JM, Merilä J. 2008. Comparative studies of quantitative trait and neutral marker divergence: a meta-analysis. J. Evol. Biol. 21:1–17.

Lemos B, Meiklejohn CD, Cáceres M, Hartl DL. 2005. Rates of divergence in gene expression profiles of primates, mice, and flies: stabilizing selection and variability among functional categories. Evolution. 59:126–137.

Levy SF, Siegal ML. 2008. Network hubs buffer environmental variation in Saccharomyces cerevisiae. PLoS Biol. 6:2588–2604.

Li H, Handsaker B, Wysoker A, Fennell T, Ruan J, Homer N, Marth G, Abecasis G, Durbin R. 2009. The Sequence Alignment/Map format and SAMtools. Bioinformatics 25:2078–2079.

Li W, Godzik A. 2006. Cd-hit: A fast program for clustering and comparing large sets of protein or nucleotide sequences. Bioinformatics 22:1658–1659.

López-Maury L, Marguerat S, Bähler J. 2008. Tuning gene expression to changing environments: from rapid responses to evolutionary adaptation. Nat. Rev. Genet. 9:583–593.

Merilä J. 2012. Evolution in response to climate change: in pursuit of the missing evidence. BioEssays 34:811–818.

Merilä J, Hendry AP. 2014. Climate change, adaptation, and phenotypic plasticity: the problem and the evidence. Evol. Appl. 7:1–14.

Messer PW, Ellner SP, Hairston NG. 2016. Can Population Genetics Adapt to Rapid Evolution? Trends Genet. 32:408–418.

Morris MRJ, Rogers SM. 2013. Overcoming maladaptive plasticity through plastic compensation. Curr. Zool. 59:526–536.

Murren CJ, Auld JR, Callahan H, Ghalambor CK, Handelsman CA, Heskel MA, Kingsolver JG, Maclean HJ, Masel J, Maughan H, et al. 2015. Constraints on the evolution of phenotypic plasticity: limits and costs of phenotype and plasticity. Heredity. 115:293–301.

Narum SR, Campbell NR, Meyer KA, Miller MR, Hardy RW. 2013. Thermal adaptation and acclimation of ectotherms from differing aquatic climates. Mol. Ecol. 22:3090–3097.

Nei M, Suzuki Y, Nozawa M. 2010. The Neutral Theory of Molecular Evolution in the Genomic Era. Annu. Rev. Genomics Hum. Genet. 11:265–289.

O’Hara RB, Merilä J. 2005. Bias and precision in QST estimates: Problems and some solutions. Genetics 171:1331–1339.

Papakostas S, Vøllestad LA, Bruneaux M, Aykanat T, Vanoverbeke J, Ning M, Primmer CR, Leder EH. 2014. Gene pleiotropy constrains gene expression changes in fish adapted to different thermal conditions. Nat. Commun. 5:4071.

Parter M, Kashtan N, Alon U. 2007. Environmental variability and modularity of bacterial metabolic networks. BMC Evol. Biol. 7:169.

Pearson JC, Lemons D, McGinnis W. 2005. Modulating Hox gene functions during animal body patterning. Nat. Rev. Genet. 6:893–904.

Phifer-Rixey M, Bomhoff M, Nachman MW. 2014. Genome-wide patterns of differentiation among house mouse subspecies. Genetics 198:283–297.

Pigliucci M. 2006. Phenotypic plasticity and evolution by genetic assimilation. J. Exp. Biol. 209:2362–2367.

Price TD, Qvarnstrom A, Irwin DE. 2003. The role of phenotypic plasticity in driving genetic evolution. Proc. R. Soc. B Biol. Sci. 270:1433–1440.

Puebla O, Bermingham E, McMillan WO. 2014. Genomic atolls of differentiation in coral reef fishes (Hypoplectrus spp., Serranidae). Mol. Ecol. 23:5291–5303.

Reusch TBH. 2014. Climate change in the oceans: evolutionary versus phenotypically plastic responses of marine animals and plants. Evol. Appl. 7:104–122.

Rifkin SA, Kim J, White KP. 2003. Evolution of gene expression in the Drosophila melanogaster subgroup. Nat. Genet. 33:138–144.

Risso D, Ngai J, Speed TP, Dudoit S. 2014. Normalization of RNA-seq data using factor analysis of control genes or samples. Nat. Biotechnol. 32:896–902.

Roberge C, Guderley H, Bernatchez L. 2007. Genomewide identification of genes under directional selection: Gene transcription QST scan in diverging atlantic salmon subpopulations. Genetics 177:1011–1022.

Roberts A, Pachter L. 2012. Streaming fragment assignment for real-time analysis of sequencing experiments. Nat. Methods 10:71–73.

Rohlfs R V., Harrigan P, Nielsen R. 2014. Modeling gene expression evolution with an extended ornstein-uhlenbeck process accounting for within-species variation. Mol. Biol. Evol. 31:201–211.

Romero IG, Ruvinsky I, Gilad Y. 2012. Comparative studies of gene expression and the evolution of gene regulation. Nat. Rev. Genet. 13:505–516.

Ruprecht C, Vaid N, Proost S, Persson S, Mutwil M, Rensing SA, al. et, Merchant SS, al. et, Chaney L, et al. 2017. Beyond Genomics: Studying Evolution with Gene Coexpression Networks. Trends Plant Sci. 22:298–307.

Salinas S, Munch SB. 2012. Thermal legacies: Transgenerational effects of temperature on growth in a vertebrate. Ecol. Lett. 15:159–163.

Schlichting CD, Wund MA. 2014. Phenotypic plasticity and epigenetic marking: An assessment of evidence for genetic accommodation. Evolution. 68:656–672.

Shama LNS, Mark FC, Strobel A, Lokmer A, John U, Mathias Wegner K. 2016. Transgenerational effects persist down the maternal line in marine sticklebacks: gene expression matches physiology in a warming ocean. Evol. Appl. 9:1096–1111.

Shaw RG, Etterson JR. 2012. Rapid climate change and the rate of adaptation: insight from experimental quantitative genetics. New Phytol. 195:752–765.

Siegal ML, Promislow DEL, Bergman A. 2007. Functional and evolutionary inference in gene networks: Does topology matter? Genetica 129:83–103.

Sikkink KL, Reynolds RM, Ituarte CM, Cresko WA, Phillips PC. 2014. Rapid Evolution of Phenotypic Plasticity and Shifting Thresholds of Genetic Assimilation in the Nematode Caenorhabditis remanei. G3Genes|Genomes||Genetics 4:1103–1112.

Smeds L, Künstner A. 2011. ConDeTri - A Content Dependent Read Trimmer for Illumina Data.Donlin MJ, editor. PLoS One 6:e26314.

Snell-Rood EC, Van Dyken JD, Cruickshank T, Wade MJ, Moczek AP. 2010. Toward a population genetic framework of developmental evolution: The costs, limits, and consequences of phenotypic plasticity. BioEssays 32:71–81.

Soyer OS, O’Malley MA. 2013. Evolutionary systems biology: what it is and why it matters. BioEssays 35:696–705.

Supek F, Bošnjak M, Škunca N, Šmuc T, Rivals I, Personnaz L, Taing L, Potier M-C, Ashburner M, Ball C, et al. 2011. REVIGO Summarizes and Visualizes Long Lists of Gene Ontology Terms.Gibas C, editor. PLoS One 6:e21800.

Szklarczyk D, Franceschini A, Wyder S, Forslund K, Heller D, Huerta-Cepas J, Simonovic M, Roth A, Santos A, Tsafou KP, et al. 2015. STRING v10: Protein-protein interaction networks, integrated over the tree of life. Nucleic Acids Res. 43:D447–D452.

Wagner GP. 1996. Homologues, Natural Kinds and the Evolution of Modularity. Am. Zool. 36:6–43.

Wagner GP, Altenberg L. 1996. Perspective?: Complex Adaptations and the Evolution of Evolvability. Evolution (N. Y). 50:967–976.

Wagner GP, Pavlicev M, Cheverud JM. 2007. The road to modularity. Nat. Rev. Genet. 8:921–931.

Walworth NG, Lee MD, Fu F-X, Hutchins DA, Webb EA. 2016. Molecular and physiological evidence of genetic assimilation to high CO 2 in the marine nitrogen fixer Trichodesmium. Proc. Natl. Acad. Sci. 113:E7367–E7374.

Weir Cockerham, B.C. 1984. Estimating F-Statistics for the Analysis of Population Structure. Evolution (N. Y). 38:1358–1370.

Whitehead A, Crawford DL. 2006. Variation within and among species in gene expression: Raw material for evolution. Mol. Ecol. 15:1197–1211.

Whitlock MC. 2008. Evolutionary inference from QST. Mol. Ecol. 17:1885–1896.

De Wit P, Pespeni MH, Palumbi SR. 2015. SNP genotyping and population genomics from expressed sequences - Current advances and future possibilities. Mol. Ecol. 24:2310–2323.

Xiong F, Ji Z, Liu Y, Zhang Y, Hu L, Yang Q, Qiu Q, Zhao L, Chen D, Tian Z, et al. 2017. Mutation in SSUH2 Causes Autosomal-Dominant Dentin Dysplasia Type I. Hum. Mutat. 38:95–104.

